# Identification of a multipotent lung progenitor for lung regeneration

**DOI:** 10.1101/2022.07.07.498730

**Authors:** Chava Rosen, Elias Shetzen, Irit Milman-Krentsis, Yuan Qi, Ran Orgad, Xiaohua Su, Raj Yadav, Michal Shemesh, Adi Biram, Ziv Shulman, Smadar Eventov-Friedman, Mukesh Maharjan, Jing Wang, Moshe Biton, Yair Reisner

**Affiliations:** Department of Immunology and Regenerative Biology, Weizmann Institute of Science, Rehovot, Israel; Department of Stem Cell Transplantation and Cellular Therapy, University of Texas MD Anderson Cancer Center, Houston, Texas, USA; Department of Neonatology, Edmond and Lily Safra Children’s Hospital, Sheba Medical Center, Israel; Cell Observatory of the MICC Life Sciences Core Facilities, Weizmann Institute of Science, Rehovot, Israel; Department of Neonatology, Hadassah Medical Center, Faculty of Medicine, Hebrew University of Jerusalem, Israel; Department of Bioinformatics and Computational Biology, MD Anderson Cancer Center, Houston, Texas, USA; Department of System Immunology

## Abstract

We recently showed that intravenous infusion of mouse or human, fetal or adult lung cells following conditioning of recipient mice leads to lung chimerism within alveolar and bronchiolar lineages, in distinct ’patches’ containing both epithelial and endothelial cells. We show here, using *R26R-Confetti* mice as donors, that these multi-lineage patches are derived from a single lung progenitor. FACS of adult mouse lung cells revealed that the putative patch-forming progenitors co-express the endothelial marker CD31 (PECAM-1) and the epithelial marker CD326 (EPCAM). Transplantation of lung cells from transgenic Cre/lox mice expressing nuclear GFP under the VEcad promoter (VEcad-Cre-nTnG), led to GFP+ patches comprising both GFP+ endothelial and epithelial cells in vivo, and in ex-vivo culture of CD326+CD31+ progenitors. Single cell RNA sequencing of CD326+CD31+ lung cells revealed a subpopulation expressing canonical epithelial and endothelial genes. Such double positive GFP+NKX2.1+SOX17+ cells were also detected by immunohistological staining in lungs of VEcad-Cre-nTnG (expressing nuclear GFP) mice in proximity to blood vessels. These findings provide new insights on lung progenitors and lung development and suggest a potential novel approach for lung regeneration.

**Summary:** We show in the present study, that multi-lineage regenerative patches in our transplantation model are derived from a single lung progenitor, co-expressing the endothelial marker CD31 and the epithelial marker CD326. These findings provide new insights on lung progenitors and lung development.

## Introduction

End-stage respiratory diseases are among the leading causes of death worldwide, with more than 5.5 million deaths annually (World Health Organization data for 2020). Today, the only potential curative treatment for these conditions is by replacement of the damaged organ with a lung transplant. Due to a shortage of suitable organs, many patients die on the transplant waiting list, and therefore lung diseases are prime candidates for stem cell therapy.

Various cell populations have been shown to exhibit regenerative potential, including lung-derived p63+ cells (1), LNEP (lineage-negative epithelial progenitors) (2) and mouse and human Sox9+ cells (3) (4). Recently, we showed that fetal lung progenitors could offer an attractive source for transplantation in mice, provided that the lung stem cell niche in the recipient is vacated of endogenous lung progenitors by adequate conditioning (5). In a procedure akin to bone marrow transplantation (BMT), a single cell suspension of mouse or human fetal lung cells harvested at the canalicular phase of gestation (20–22 weeks in human, and E15–E16 for mouse) and infused intravenously (I.V) following conditioning of recipient mice with Naphthalene (NA) and 6Gy total body irradiation (TBI), led to marked lung chimerism, in distinct ’patches’ containing both epithelial and endothelial cells. This chimerism was associated with significantly improved lung function (5). More recently, this approach for lung chimerism induction was extended to transplantation of a single cell suspension of adult mouse lung donors (6), (7), (8), requiring about three-fold higher cell doses to attain a similar level of chimerism (6).

Considering that the lung patches are observed within confined borders similarly to spleen colonies known to be derived from a single multi-potent hematopoietic progenitor, we hypothesized that these lung patches might represent a clonal derivation from a yet unknown multi-potent lung progenitor. To address this hypothesis, we transplanted fetal or adult lung cells from *Rosa26-Confetti* mice bearing a multi-color Cre reporter system (9). This four-color Cre recombination system provides a fetal or adult lung cell preparation in which each cell expresses just one randomly determined colour. Thus, the likelihood that each cell within a doublet in the transplanted cell population would be of the same colour is markedly reduced (10). We demonstrate that all donor-derived lung patches developing after transplantation are monochromatic, strongly supporting the clonal origin of donor-derived lung patches observed after transplantation, in striking resemblance to the spleen colony-forming cells typically identified after BMT. These results provide definitive evidence of a single multi-potential lung progenitor capable of differentiating into diverse lung cell lineages.

In line with our observation that a large proportion of the patches contain both endothelial and epithelial cells and that each such patch is derived from a single progenitor, we searched for a putative multipotent lung progenitor capable of differentiating along these two distinct lineages. Further characterization of these progenitor cells by FACS showed that they exhibit a dual profile, expressing markers typical of both endothelial and epithelial fates. Furthermore, the presence of a unique cluster of double-positive cells is supported by single cell RNA-sequencing (scRNA-seq) and by immunohistology of lungs of VEcad-Cre-nTnG transgenic mice. Apart from its potential for translational studies aiming at the correction of different lung diseases, the Identification of such progenitors could also contribute to basic studies aimed at a better understanding of fetal lung development, as well as the understanding of steady-state maintenance of different cellular lineages in the adult lung.

## Results

### Long term chimerism in recipient lungs after transplantation of td-Tomato-labeled cells

In our previous studies we demonstrated by immunohistology and by quantitative morphometric analysis robust lung chimerism at 2-4 months after transplantation of fetal or adult lung cells into mice conditioned with NA and 6Gy TBI (5), (6). We also demonstrated in these chimeric mice significant functional benefit as measured by *Flexivent.* We have now extended the chimerism follow-up period to 6-8 months and found a marked level of donor-derived patches (Fig. 1A and Supplementary Fig. 1). This high level of donor-type chimerism was also documented by FACS analysis of a cell suspension of dissociated chimeric lungs, showing an average of 35 ±18 % donor-derived cells (range= 9% to 64%) within the single live CD45- cells (Fig. 1B, C). Further immunohistological analysis of the donor-derived patches revealed extensive numbers of alveoli and bronchial structures (Fig. 1D-G). The former was initially indicated by triple staining for TdTomato, E-cadherin (Ecad), and CD31 (Fig. 1D) and the latter by triple staining for TdTomato, Nkx 2.1 and Ecad (Fig. 1E), TdTomato, cytokeratin (CK) and Ecad (Fig. 1F) or TdTomato, CC10+, and CD31 (Fig. 1G).

**Fig. 1.**
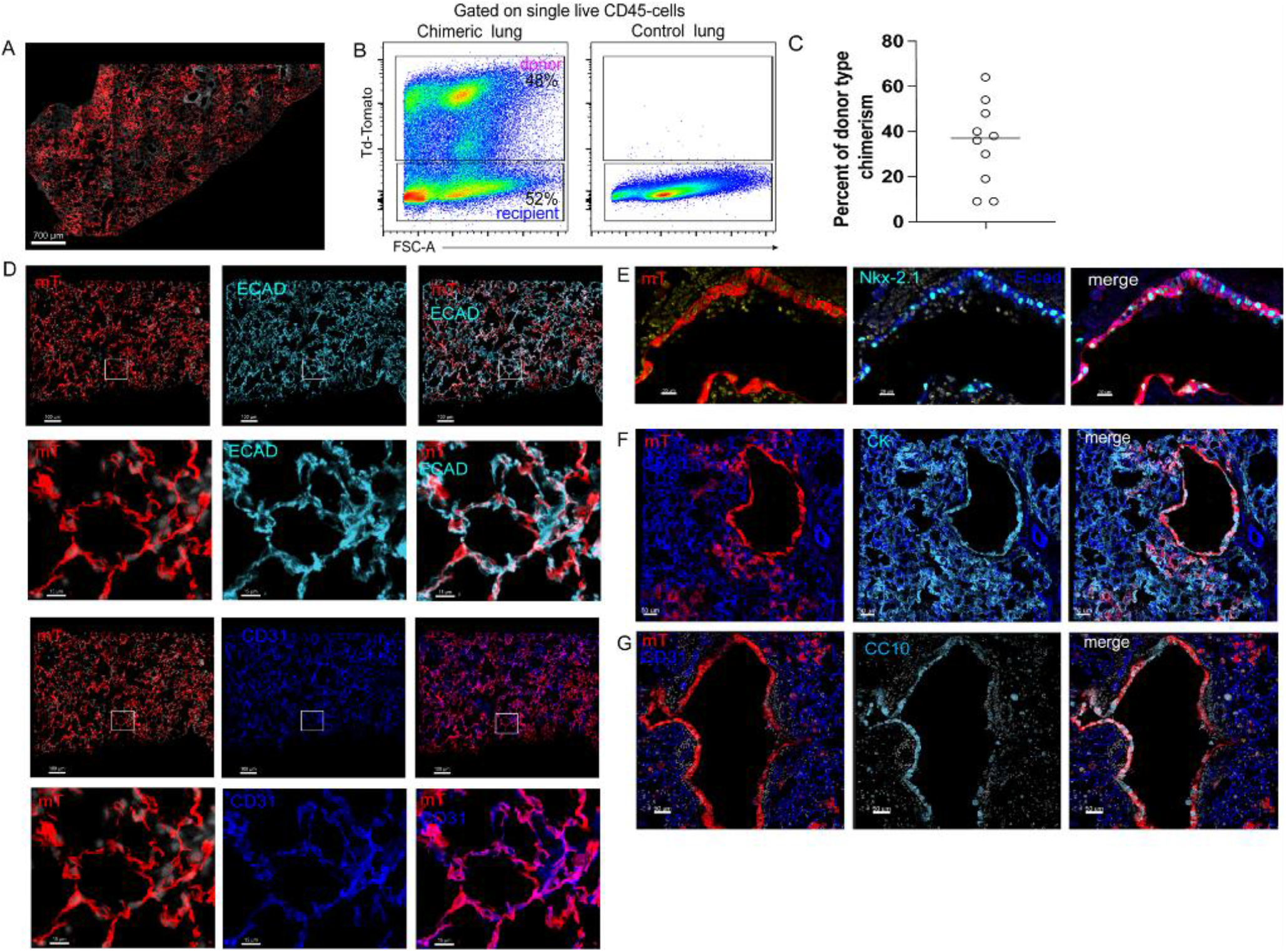
Staining of epithelial and endothelial cells within donor-derived regenerative patches in the chimeric lung at 6-8 months post-transplantation. (A) Representative image of chimeric lung, demonstrating extensive chimerism and robust donor derived TdTomato (mT-membranous tomato) engrafted regions under low magnification. Scale bar=700um. (B) Representative FACS analysis of chimeric lung compared to non-chimeric lung, demonstrating 48 % of donor-type chimerism. (C) Percentage of donor type chimerism, evaluated by FACS of *n*=10 mice from two independent experiments. The whole mount images of the lungs of these mice are shown in Supplementary Fig. 1. (D) Staining of the chimeric lung by anti-E-cadherin (cyan) and anti-CD31 (blue) antibodies, demonstrating the presence of epithelial and endothelial compartments in the chimeric lung. Top images show staining for E-cadherin, scale bar=100 um, top middle image is a close-up image of the boxed area, scale bar=15um. Middle bottom image shows staining of the same region for CD31, scale bar=100um. bottom image is a close-up of the boxed area, scale bar=15um. (E) Staining of donor-derived TdTomato+ bronchial patch with anti-Ecad (membranous, blue) and anti-Nkx-2.1 antibody (nuclear, cyan) and Hoechst staining of nuclei (yellow); scale bar=20um. (F) Staining of donor-derived TdTomato+ broncho-alveolar patch with anti-CD31 (blue), and anti-cytokeratin (cyan) antibodies; scale bar=50um. (G) Staining of donor derived TdTomato+ bronchial patch with anti-CD31(blue) and anti-CC10 (cyan) antibodies and Hoechst dye for nuclei (grey); scale bar=50um. The images shown are representative of *n*=3 mice.

Further analysis of the donor-derived alveolar structures within the patches shows marked levels of AT1 and AT2 cells. Donor-derived AT1 cells were depicted by staining for TdTomato and HOPX, TdTomato and anti-AQP-5 (Fig. 2A, B), or by TdTomato and staining of single molecule fluorescence in situ hybridization (smFISH) stain for AQP-5 (Fig. 2C), while donor-derived endothelial cells within the patches were clearly documented by staining with nuclear anti-ERG antibody (Fig. 2D). AT2 cells were identified by triple staining for TdTomato, NKX2.1 and LAMP3 (Fig. 3A), by TdTomato and ProSPC (Fig. 3B) or by TdTomato and smFISH for SPC (Fig. 3C, D).

**Fig. 2.**
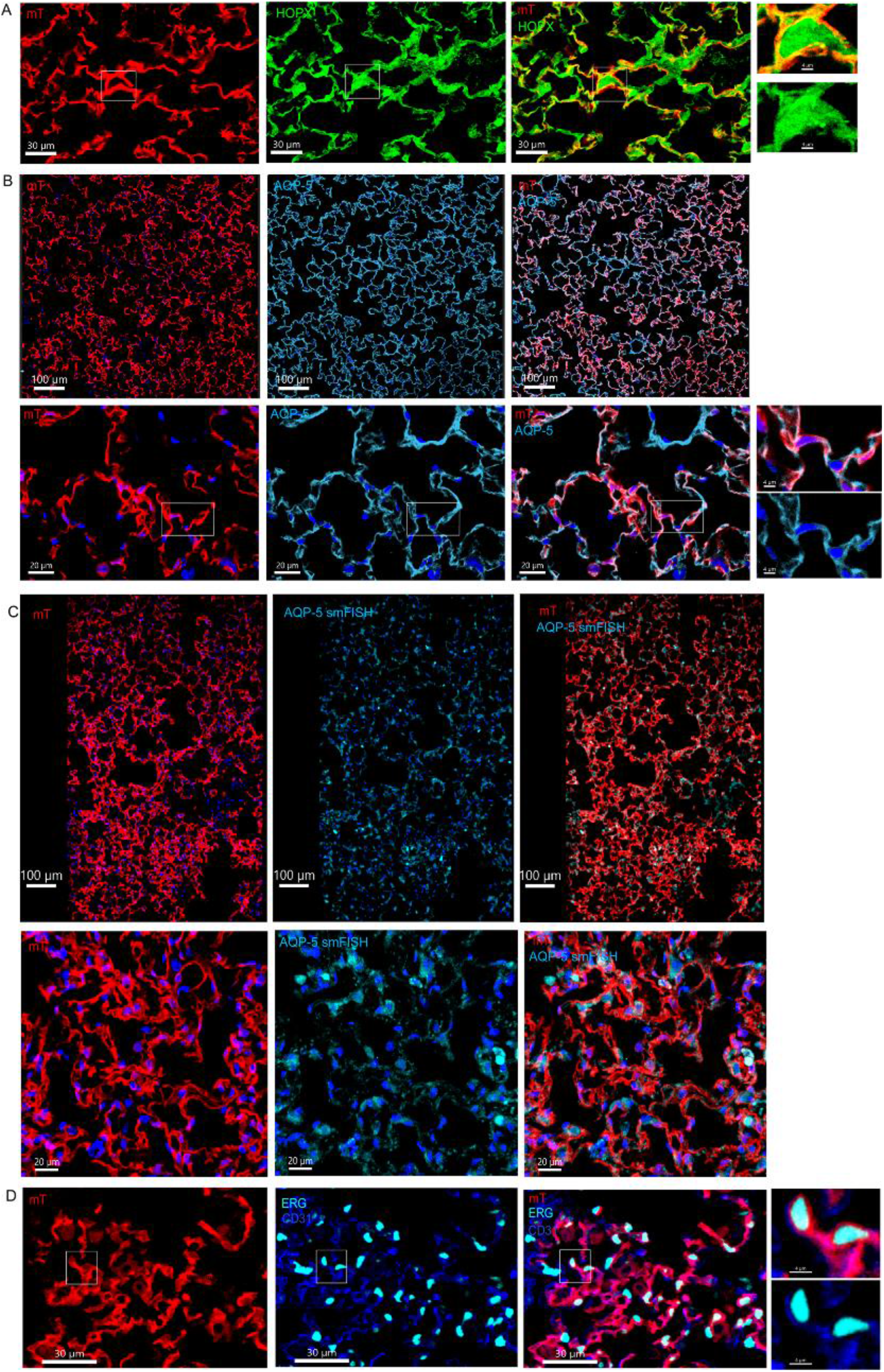
Staining of AT1 and endothelial cells within donor-derived regenerative patches in the chimeric lung at 6-8 months post-transplantation. Staining of TdTomato+ cells with anti-HOPX (A, green) or anti-AQP-5 (B, cyan) antibodies, (C) FISH Staining of TdTomato+ cells with AQP-5 RNA probe. (D) Staining of TdTomato+ cells with anti-ERG (cyan) and anti-CD31 (blue) antibodies. The images are representative of *n*=3 chimeric mice.

**Fig. 3.**
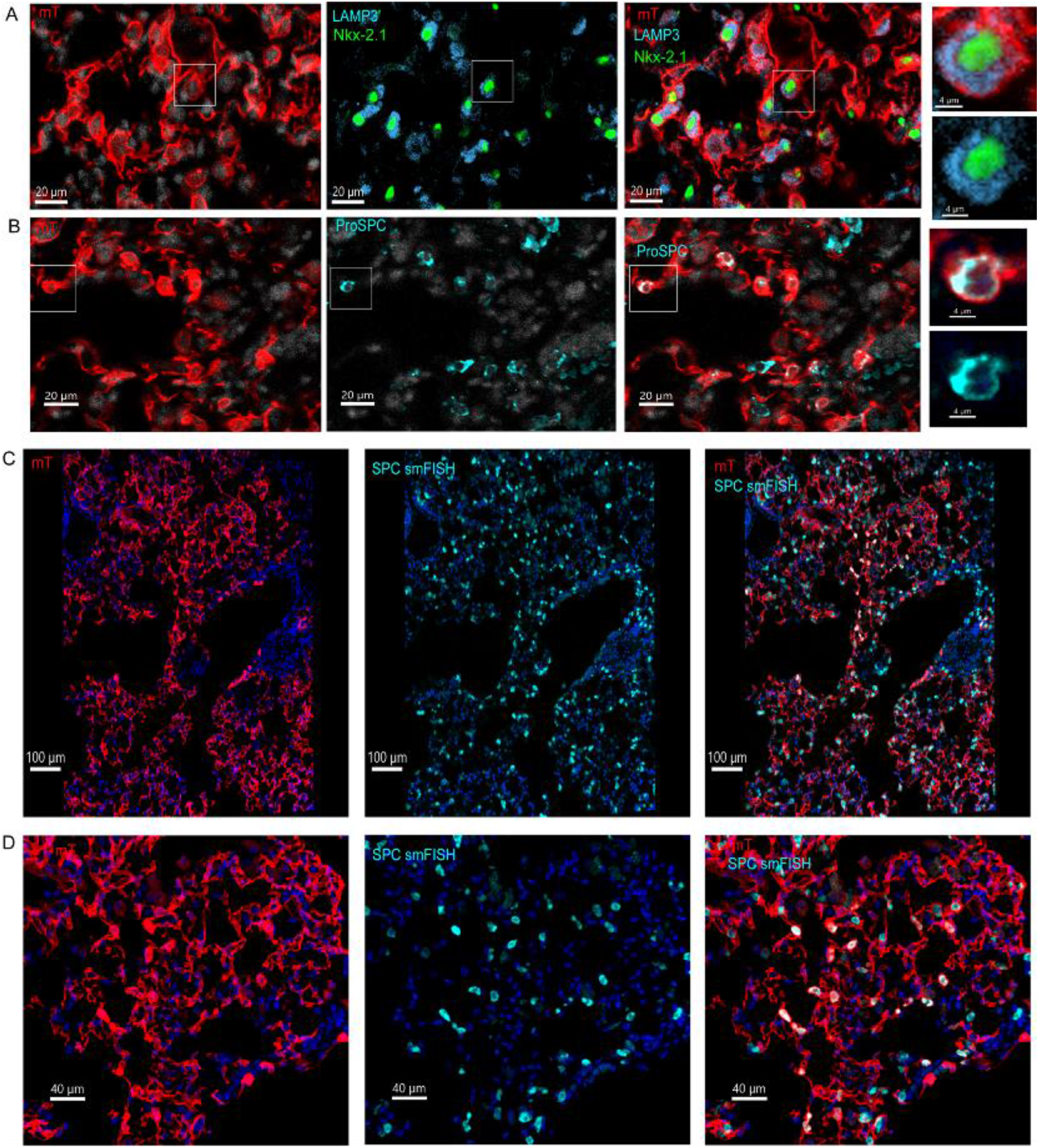
Staining of AT2 within donor-derived regenerative patches in the chimeric lung at 6-8 months post-transplantation. (A) Staining of TdTomato+ cells with anti-LAMP-3 (cyan) and anti-Nkx 2.1(green) antibodies. (B) Staining of TdTomato+ cells with anti-ProSPC (cyan) antibody. Scale bar= 20um. (C, D) FISH Staining of TdTomato+ cells with SPC RNA probe (cyan) under low (C, scale bar=100 um) and higher magnification (D, scale bar =40um respectively). Images are representative of *n*=3 mice.

Notably, staining of single molecule FISH (smFISH) for AQP-5 and ProSPC as well as the anti-ERG staining, enabled to quantitate the percentage of donor-derived AT1 (47% out of all quantified AT1 cells per field), AT2 (40% out of all quantified AT2 cells per field), and endothelial cells (52% out of all quantified endothelial cells per field) in 0.2 mm^2^ of donor derived regions within chimeric lung (Supplementary Fig. 2) .

### Transplantation of lung progenitors from fetal or adult Cag-Cre ER2 ‘Confetti’ mice results in monochromatic patches

To unequivocally examine our hypothesis that the lung patches observed following transplantation are derived from a single progenitor, we carried out a set of experiments making use of the multicolor ‘*R26R-Confetti*’ mouse reporter system (9) as donors. In these mice, the recombination process is independent in each cell and stochastic. Consequently, all daughter cells will produce the same fluorescent protein, and if indeed all the cells within each patch are derived from a single progenitor, they should all be of the same color. Multiple studies used this system to demonstrate clonal behavior in organs such as the intestine (9),(11), lung(12), (13), brain(14), kidney, mammary gland, and others (15), (16), (17), (18), (19).

Thus, we began investigating lung patch formation following transplantation of lung cell suspension from E16 *R26R-Confetti* donors into RAG-2 recipients preconditioned with NA and 6Gy TBI (see schematic overview in Supplementary Fig. 3A and details in Methods). Tamoxifen (TMX) was administered to pregnant females at E12, the embryos were harvested at E16, and the fetal lungs containing fluorescent cells were isolated under a fluorescent microscope. Two-photon micrographs depict the typical population of monochromatic cells prior to transplantation, each expressing one of the four fluorescent tags (Supplementary Fig. 3B, C).

Transplantation of fetal lung cells from Tamoxifen-induced donors into NA-treated and irradiated recipient mice, resulted in discrete monochromatic lung patches, each expressing one of the four fluorescent proteins, strongly indicating that each patch is likely derived from a single lung progenitor (Supplementary Fig. 3D).

Next, based on our recent finding that patch-forming progenitor cells are also present in the adult mouse lung, we applied the same approach to analyze the origin of lung patches after transplantation of adult lung cells. To this end, we used the protocol described above, but since the frequency of patch-forming cells in the adult lung is about 3-4 fold lower compared to that found in E16 fetal lungs (6), and only a fraction of the cells undergo the Cre recombination, we used a higher dose (16x10^6^) of lung cells for transplantation (Fig. 4A) As shown in Fig. 4B and Supplementary Movie 1, following TMX treatment, monochromatic GFP or YFP or RFP or CFP fluorescent cells could be documented by two-photon microscopy within the adult lung (Fig. 4B), and as shown by confocal microscopy, these tagged cells were randomly distributed throughout the entire lung (Fig. 4B). To evaluate the clonality of donor-derived patches at 8 weeks after transplantation, we used confocal, two-photon, and light sheet microscopy.

**Fig. 4.**
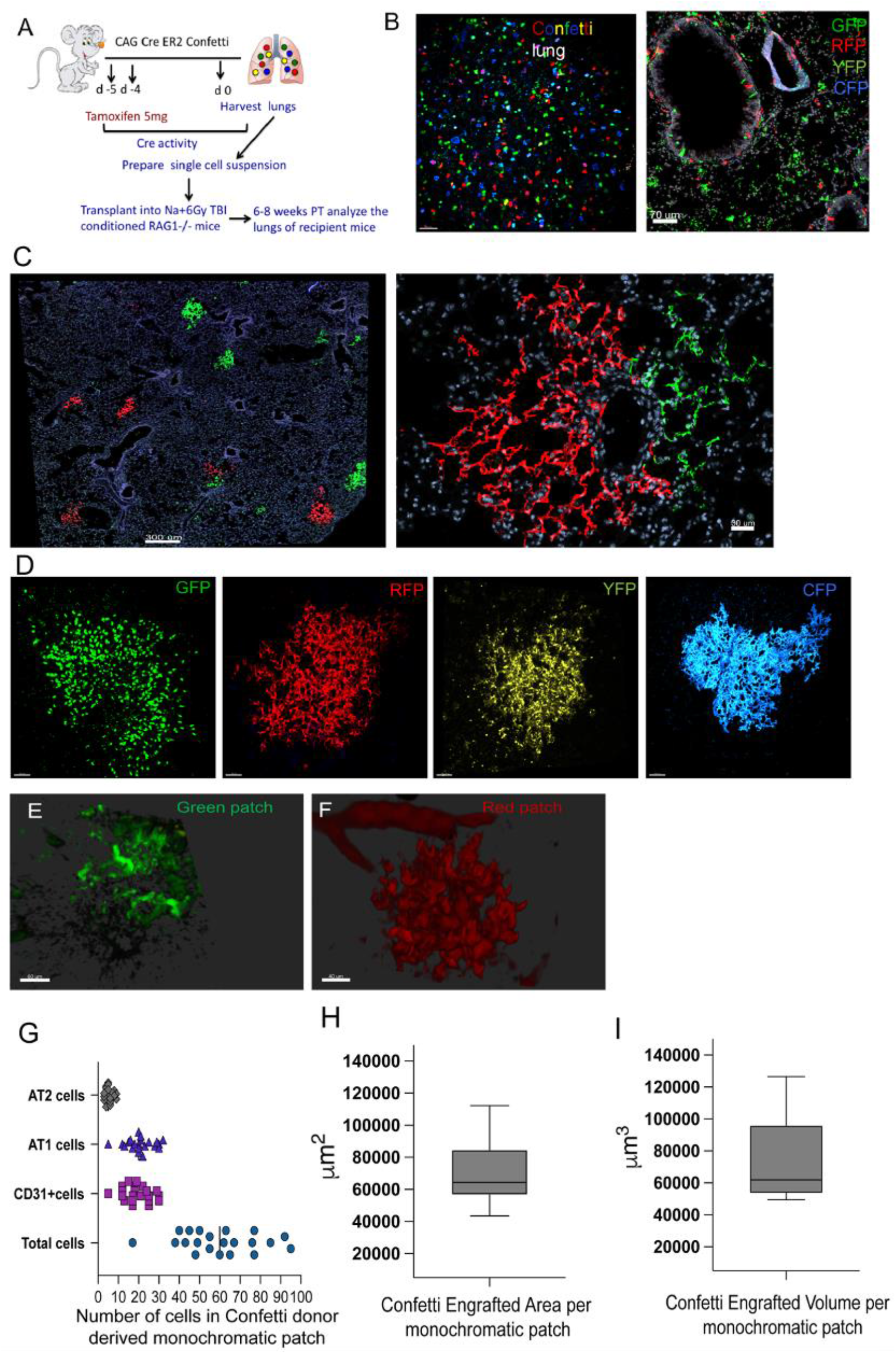
Lung chimerism analysis at 8 weeks after transplantation of adult *R26R-Confetti* lung cells. (A) Schematic presentation of the experimental procedure. (B, left image) Two-photon microscopy of the monochromatic patches of transplanted adult lungs. Single cells, each expressing one of the four tag colors after Cre recombination; *n*=3. (B, right image) Confocal image of adult *R26R-Confetti* lung slice demonstrating bronchial and alveolar parts of the lung tissue, and diffuse and random localization of the fluorescent cells within the donor lung. The results are representative of *n*=6 *R26R-Confetti* mice from 2 independent experiments. (C) Fluorescence microscopy of a section of chimeric lung transplanted with *R26R-Confetti* lung cells, demonstrating several single-color patches (left image, scale bar=200um), and a pair of neighboring single-color patches (right image, scale bar=50um). (D) Two-photon microscopy of monochromatic patches, each exhibiting a distinct color in whole mount chimeric lung; scale bar=50um; *n*=4 chimeric lungs. Each patch with alternative fluorescence channels is illustrated in Supplementary Fig.4. Transparent chimeric lung produced by a clearing procedure was analyzed by light sheet microscopy, *n*=2 chimeric lungs were evaluated by LSM; in (E) scale bar= 60um and in (F), scale bar=40um. (G-I) Quantitative composition analysis of monochromatic patches derived from transplanted Confetti donors. (G) Number of AT1, AT2, and endothelial cells per monochromatic patch, in a total of *n*=19 patches from *n*=5 chimeric mice evaluated 8 weeks after transplantation. (H-I) Box plots, depicting the area and volume occupied by each patch. Quantitative evaluation of the frequency of donor-derived AT1(AQP-5+ or HOPX+), AT2 (SPC) or endothelial (CD31+) cells within each patch was performed on serial slices from different mice (*n*=5) using manual evaluation as well as Imaris software. The entire dataset, including the patch volume, number of nuclei (Imaris “surface” algorithm), and number of epithelial and endothelial cells (calculated manually), was collected from a total of 19 monochromatic patches.

Evaluation of chimeric lung slices by fluorescence microscopy, allowing for the detection of two chromophores (red and green) demonstrated discrete monochromatic patches (Fig. 4C). Two-photon snapshots of whole mount chimeric lung detected all 4 chromophores, with monochromatic patches expressing either membranous CFP, nuclear GFP, or cytoplasmatic RFP or YFP (Fig. 4D). Each monochromatic patch tested under the alternative fluorescence channels are shown in Supplementary Fig. 4. Notably, taken together we found that in 50 different fields from 15 chimeric mice (12 transplanted with adult lung cells and 3 with fetal cells), all the patches are monochromatic, which strongly negates the possibility that these patches are derived from two different cells. The full depth two-photon microscopy scan of these monochromatic patches in the whole mount lung tissue is presented in Supplementary Movie 2.

Thus, testing our experimental distribution of single-color (n=50) vs. two-color (n=0) clones against the theoretical distribution of 25% vs. 75% (χ21 = 150), by χ2 distribution test, suggests that it is highly unlikely that any clone is derived from two cells (p < 0.001). In addition, as shown in Fig. 4E, F, and Supplementary Movie 3, light sheet microscopy of “cleared” chimeric lungs also revealed distinct monochromatic patches. Importantly, double staining analyzed by confocal microscopy shows that each monochromatic patch exhibits both epithelial and endothelial cells (Supplementary Fig. 5-7).

Notably, most of the patches contained both donor-derived epithelial and endothelial cells. Thus, the average number of donor-derived AT1 and AT2 alveolar cells and endothelial cells within the patches 8 weeks post-transplantation was 21±5, 6±3 and 19±5 respectively, out of 62±17 total donor-derived cells per patch Supplementary Fig. 8 and Supplementary Table 1). These results quantitatively demonstrate that each monochromatic patch contains both endothelial and epithelial cells, suggesting that a plastic multi-potent lung progenitor within fetal and adult mouse lungs can differentiate into these distinct lung lineages

### Two distinct patch-forming lung cell progenitors

Considering that a large proportion of the observed patches in chimeric lungs contain both endothelial and epithelial cells (5) (Fig. 1-3) and that each such patch is derived from a single progenitor, we searched for a putative multipotent lung progenitor capable of differentiating along these two distinct lineages. As shown in Fig. 5A, FACS analysis of the CD45- lung cell population for expression of epithelial (CD326+) and endothelial (CD31+) markers clearly reveals the presence of a unique CD45-CD326+CD31+ double-positive subpopulation exhibiting a wide range of CD326+ expression. This analysis revealed in 40 mice from 23 independent experiments, an average percentage of 91±5.5 (range 70-97%) single cells, 35±24.2% of CD45- (range 9-72%) cells and 8.2±2.7% (range 4.5-13.3%) CD45-CD326+CD31- cells; 3±1.5% CD45-CD326+CD31+ cells (range1-5.8%); 51±9.9% CD45-CD326-CD31+ cells (range 26-63%); and 26±6.5% CD45-CD326-CD31- cells (range 23-41%) (Fig. 5B). Of note, the percentage of CD326+ cells, is relatively low in FACS assays due to the low retrieval of these cells by the enzymatic lung digestion. Next, we attempted to isolate by FACS the four sub-populations, as shown schematically in Fig. 5C, and define their capacity to form patches. To trace donor-derived patches 8 weeks after transplantation of purified cell populations, we made use of labeled donor mice, including GFP (C57BL/6-Tg (CAG-EGFP)1Osb/J), mTmG (*Gt(ROSA)26Sor^tm4(ACTB-tdTomato,-EGFP)Luo^*/J), and nTnG *(Gt(ROSA)26Sor^tm1(CAG-tdTomato*,-EGFP*)Ees^*/J) mice (mTmG mice express membrane TdTomato, and nTnG mice express nuclear TdTomato). The unique localization of the fluorescent tag either in the membrane or the nucleus of the sorted lung cells enabled tracking of membrane-bound, cytoplasmatic and nuclear epithelial and endothelial markers at the single cell level within the monochromatic patches of chimeric lungs, and to characterize the cellular composition of the patches after transplantation of the sorted lung cell subpopulations.

**Fig. 5.**
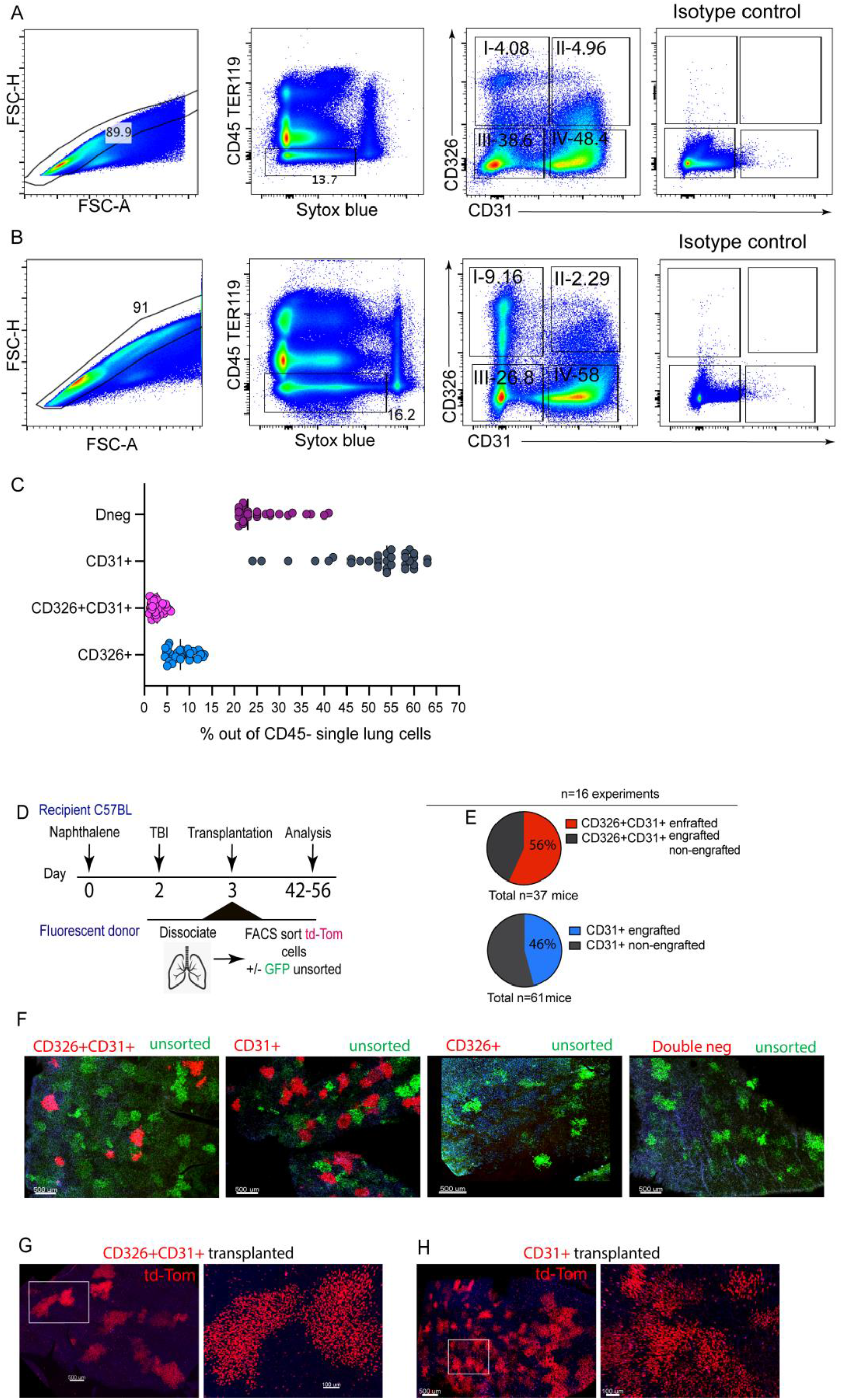
Donor-derived lung patches after transplantation of different lung cell sub-populations. (A, B) Two different examples of FACS analysis of CD45- lung cell subpopulations prior to transplantation with different percentages of CD326+CD31-, CD326+CD31+, CD326-CD31+, and CD326-CD31- cells, used in transplantation experiments and FACS strategy of lung cells: Lung cells were first gated on single cells based on forward scatter area and height, and thereafter on CD45- TER119- live cells (Sytox blue negative). The gated cells were then sorted into four fractions: I) CD326+CD31-; II) CD326+CD31+; III) CD326-CD31-; IV) CD326-CD31+. (C) Percentage of each subpopulation for *n*=40 mice from 23 independent experiments, including 16 transplantation experiments out of the CD45- non-hematopoietic lung cell population. (D) Schematic presentation of the transplantation experiments using FACS-purified cells. (E) Percentage of chimeric mice exhibiting donor-derived patches out of the total number of transplanted mice (n=16 experiments). (F) Donor-derived lung patches 6 weeks after transplantation of 0.3-0.5x10^6^ sorted double positive CD326+CD31+ cells, or single positive CD326-CD31+ endothelial cells from nTnG donors mixed with 0.5x10^6^ unsorted cells from GFP+ donors (green), as compared to CD326-CD31+ and CD326-CD31- that did not form regenerative patches. The whole mount confocal images shown for each group are representative of *n*=16 experiments, with at least *n*=10 mice in each group; scale bar=500um. Out of the total transplanted mice (n=98), 70% were transplanted with sorted TdTomato+ and unsorted GFP cells, and 30% were transplanted only with the FACS purified TdTomato+ cells. (G, H) Representative images of whole mount lungs of mice transplanted with 0.5X10^6^ nTnG CD326+CD31+ or CD31+cells, without GFP+ co-transplanted unsorted lung cells. Donor-derived red patches express nuclear TdTomato. Images under low (scale bar=500 um) and high magnification (scale bar=100um) are representative of *n*=5 mice in each group.

Notably, in a total of 16 experiments transplanting sorted cells, patch-forming activity could be found only upon transplantation of CD45-CD326+CD31+ double positive (21 of 37 mice) or single positive CD45-CD326-CD31+ cells (28 out of 61 mice) (Fig. 5D, E). No patch-forming activity was found in our transplantation model in the single positive CD45-CD326+ CD31-epithelial cell fraction, or in the triple negative CD45-CD326-CD31- population (Fig. 5F). It is possible that sorted cells might need additional facilitating cells in the donor cell preparation for the early steps of colonization in the recipient’s lungs. We therefore attempted to transplant 0.3-0.5 x10^6^ sorted cells from lungs of mTmG or nTnG mice, together with (Fig. 5F) or without (Fig. 5G, H) 0.5x10^6^ unsorted lung cells from GFP+ donors. These experiments clearly demonstrate that transplanted FACS-purified CD326+CD31+ and CD326-CD31+ cells are capable of forming patches even in the absence of potentially supporting cells from the co-transplanted unseparated GFP+ lung cell preparation (Fig. 5G, H), while CD326+CD31-single positive epithelial cells are unable to form patches even in the presence of unseparated GFP+ cells.

Staining of the donor-derived patches for epithelial alveolar markers including AQP-5, HOPX and SPC, or for endothelial markers including CD31, ERG, and SOX17, demonstrated that the patches formed following transplantation of the single positive CD326-CD31+ endothelial cells were comprised predominantly of endothelial cells (Fig. 6A, B), while patches formed after transplantation of the double positive CD326+CD31+ subpopulation contained both endothelial and epithelial cells (Fig. 6C-G). This result is strongly supported by the use of nuclear staining to distinguish cellular borders between neighboring cells. To this end, co-localization of the donor-derived nT (nuclear tomato) marker with nuclear ERG/SOX17 or with expression of AT1 hallmark marker HOPX enabled us to identify donor-derived endothelial or AT1 epithelial cells, respectively (Fig. 6C-E). A similar distinction of AT2 cells could be attained by staining with anti-SPC antibody (Fig. 6F) or a single molecule RNA (smRNA) FISH probe for SPC (Fig. 6G). Quantitative analysis of the cell numbers comprising the patches suggests that those found after transplantation of CD326+CD31+ lung cells were larger than those found after transplantation of CD326-CD31+ cells (Fig. 6H). Taken together, this analysis revealed a difference in cellular composition between the two types of patches, with co-localization of endothelial and epithelial cells mostly found in the patches formed following transplantation of the double positive CD326+CD31+ lung progenitors (Fig. 6I).

**Fig. 6.**
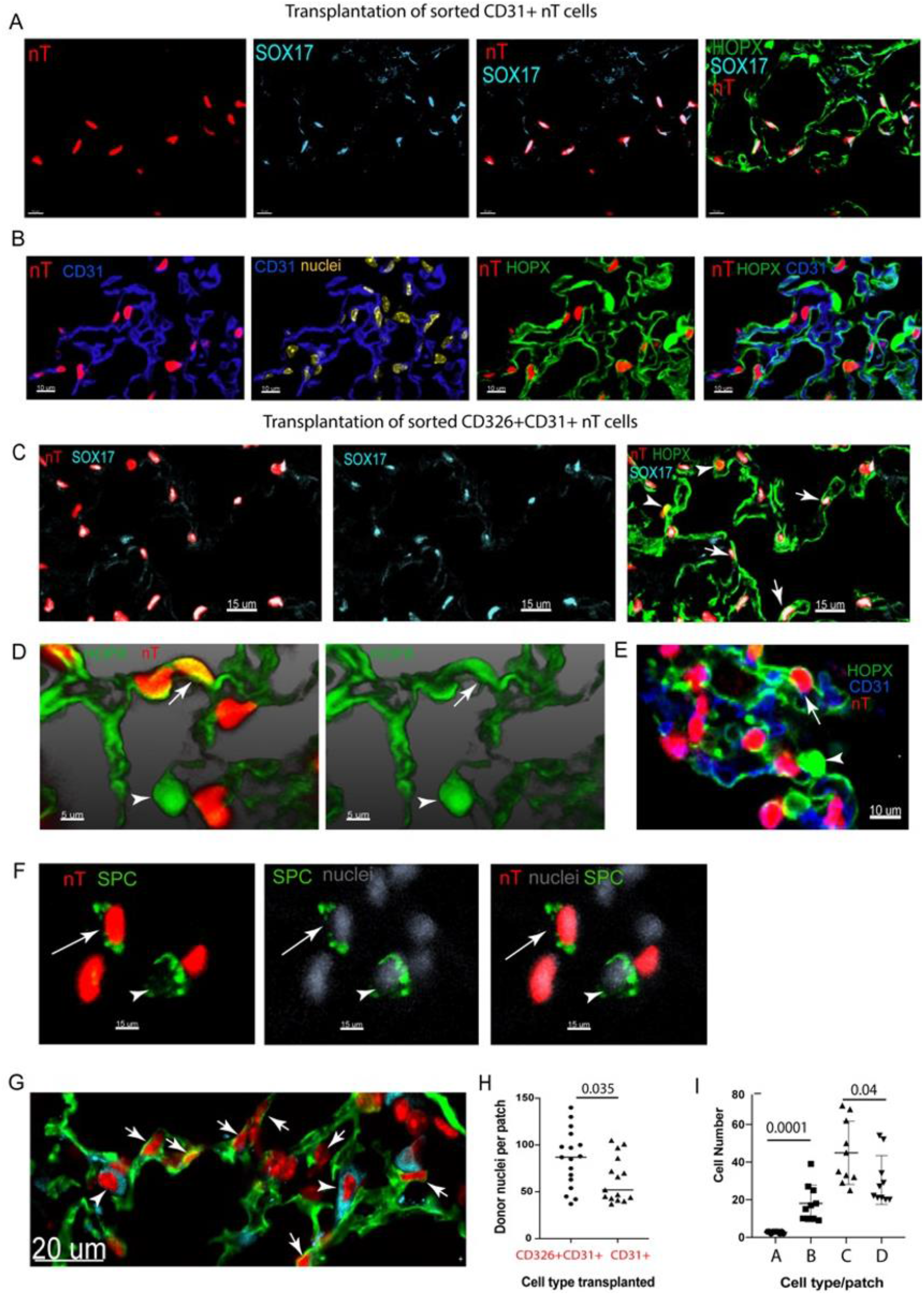
Different cellular composition of donor-derived patches after transplantation of sorted CD326-CD31+ versus CD326+CD31+ lung cell subpopulations. (A) Staining of typical lung patches derived from sorted nTnG CD326-CD31+ lung cells (red). Red nuclei positive for the endothelial nuclear marker SOX17(cyan) indicate donor-derived endothelial cells; scale bar=10um. (B) Staining for cell surface endothelial marker CD31(blue) and epithelial marker HOPX (green) in donor-derived patches formed after transplantation of CD326-CD31+ cells. Left and middle left - staining for nTnG (red) and CD31 (blue) indicating donor-derived endothelial cells. Middle right -Staining for HOPX; right – triple staining for nTnG+ (red), CD31 (blue) and HOPX (green). Donor-derived endothelial CD31+ cells reside in close proximity to recipient HOPX+ AT1 cells (indicated with arrowheads), while donor-derived AT1 cells were not detected; scale bar=10um. (C-F) Typical staining of lung patches derived following transplantation of sorted nTnG CD326+CD31+ cells, showing donor-derived epithelial and endothelial cells. (C) Left: double staining for nTnG (red) and SOX17 (cyan); Middle: single staining for SOX17; Right: Triple staining for nTnG (red), SOX17 (cyan) and HOPX (green), showing donor derived AT1 epithelial cells (arrowheads) and endothelial cells (arrows); scale bar =15um. (D) High magnification image of typical staining for HOPX showing donor (red, indicated with arrow) and host (indicated with arrowhead) AT1 cells. (E) High magnification of triple staining of CD326+CD31+-derived patch, demonstrating the presence of donor-derived (red) AT1 and endothelial CD31+ cells, in close proximity. Arrow indicates donor-derived AT1 cells, and arrowhead indicates the host-derived AT1 cells; scale bar=10um in (D) and (E). Images are representative of *n*=3 mice. (F) Staining of a donor-derived patch with anti-SPC antibody (green), arrow indicates the donor-derived nT-AT2 cell, and arrowhead shows the host AT2 cell. (G) Staining of a typical donor-derived patch for CD31 (green) and single molecule RNA FISH probe for SPC (cyan), demonstrating endothelial and AT2 cells; scale bar=20um. (H, I) Graphical summary of quantitative differences between the composition of patches derived from transplantation of CD326+CD31+ and CD326- CD31+ cells. (H) Donor-derived nuclei/patch, showing that CD326+CD31+ progenitors produce larger patches compared to CD326- CD31+ cells; p=0.035, Student’s t-test; *n*=20 patches were evaluated from *n*=3 mice in each group. (I) Absolute number of donor-derived epithelial and endothelial cells per patch, after transplantation from CD326+CD31+ or CD326-CD31+ cells: A and B -depict epithelial cells derived from transplanted CD326-CD31+ and CD326+CD31+ populations, respectively; p=0.0001, Student’s t-test. C and D depict endothelial cells derived from transplanted CD326-CD31+ and CD326+CD31+ populations, respectively; p=0.04, Student’s t-test; *n*=10 patches from 2 mice were evaluated for each group.

### Single cell RNA sequencing analysis of sorted CD326+CD31+ lung cells reveals a unique cluster expressing epithelial and endothelial canonical genes

While the sorting experiments clearly suggest that the formation of the patches, which include high levels of donor-derived epithelial and endothelial cells, originate from the double positive CD326+CD31+ cell fraction, the FACS data shown in Fig. 5A indicated a wide spectrum of CD326 positivity ranging from dim to bright cells. To further analyze the cell heterogeneity of the entire sorted CD45-CD31+CD326+ lung cell population used in our transplantation assay, and to further interrogate the potential expression of different canonical epithelial and endothelial genes within this broad CD326+CD31+ double-positive population, we used scRNA-Seq analysis of both single positive (*n*=8151 CD326+, *n*=8703 CD31+) or double positive (*n*=7689) sorted populations (Fig. 7 and Supplementary Fig. 9, *n*=4 mice). As shown in Fig. 7A, the double-positive CD45-CD31+CD326+ cell population, consists of 15 clusters (See Methods and Supplementary Table 2), and the percentage of each subset is depicted in Supplementary Fig. 9A. Doublet finder analysis (See **Methods**) shows a high rate of doublets in clusters 3, 8, and 11 (Supplementary Fig. 9B), and these clusters were not included for further analysis. Differential expression genes are provided (Supplementary Table 2), and a heatmap of the top genes in each cluster (Fig. 7B) or a dot plot of canonical epithelial and endothelial gene markers expression in the different clusters are shown (Fig. 7C). Notably, both the heatmap and the dot plot indicate that only cluster 5 exhibits cells with a dual phenotype, expressing major canonical epithelial and endothelial genes. As shown in Supplementary Fig. 9B, only 0.3% of the cells within this cluster are predicted as doublets, thus strongly negating the possibility that the dual characteristics of cluster 5 might reflect a cluster of doublets. Cluster 5 represents 9.2 % of the CD45-CD326+CD31+ cell fraction, not including clusters 3, 8 and 11, which showed a high rate of doublets (Supplementary Fig. 9B). While clusters 4 and 5 exhibited marked expression of epithelial genes such as RTKN2, AGER, AQP5, cluster 5 differed from cluster 4 in its additional expression of endothelial genes such as EPAS1, IFITM3, HPGD and Tmem (Fig. 7B). Additional well known canonical endothelial genes such as PECAM and ERG were also found to be expressed in higher levels in cluster 5 compared to cluster 4. This unique pattern of cluster 5 is also illustrated by the dot plot in Fig. 7C depicting dual expression in cluster 5 of canonical epithelial (EPCAM, Cadh1, NKX2.1, AGER) and endothelial (PECAM, ERG, SOX17, Cadh5) genes. In contrast, sorted single positive CD45-CD326+CD31-epithelial cells (Supplementary Fig. 9C and Supplementary Table 3), and sorted single positive CD45-CD326-CD31+ endothelial cells (Supplementary Fig. 9D and Supplementary Table 4) do not exhibit clusters expressing both epithelial and endothelial genes. Further analysis of cluster 5 shows a continuum of cellular heterogeneity, resulting in three subsets, with a varied expression of the endothelial and epithelial gene markers, ranging from a subset with high expression of epithelial and low endothelial markers (cluster 0), low expression of both lineages (cluster 1) and a cluster (cluster 2) with high expression of endothelial and low expression of epithelial markers (Fig. 7D and E). Notably, pseudotime analysis (See **Methods**) of cluster 5, predicted the continuum of differentiation from cluster 1 towards the two further defined lineages (Fig. 7F-H), expressing to higher extant either epithelial (cluster 0) or endothelial (cluster 2) markers (Fig. 7E). These computational analyses support the *in vivo* transplantation results and identify a small CD45-CD326+ CD31+ cell subset that may contribute to the epithelial and endothelial lineages of the lung.

**Fig. 7.**
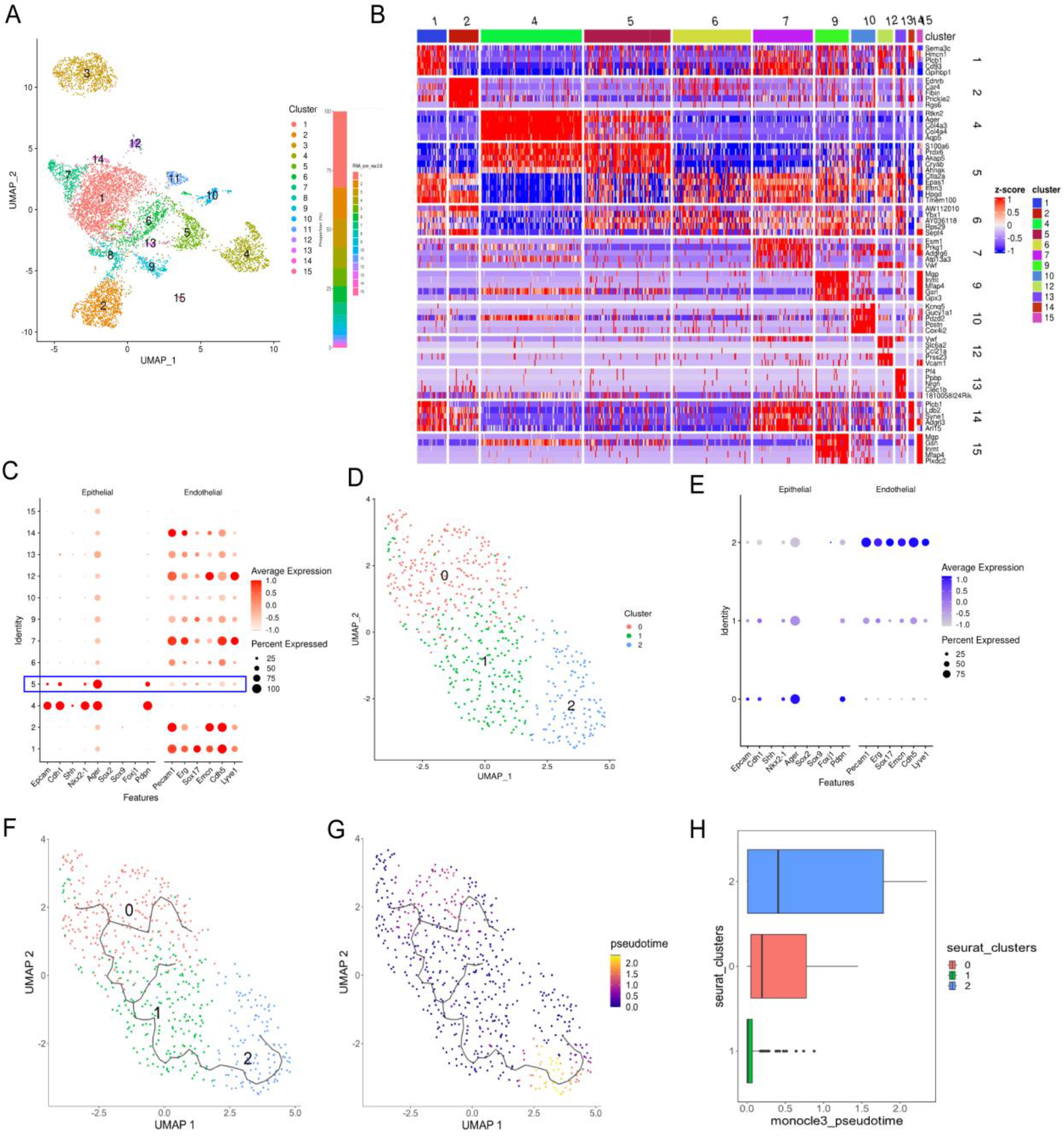
Single cell RNA transcriptome analysis of FACS purified CD326+ CD31+ cells. (A) UMAP visualization of 7689 CD326+ CD31+ cells, where individual points correspond to single cells (See Methods). Cells are colored by their assignment to clusters (*n*=4 mice). (B) CD326+CD31+ cell-subset signatures. The relative expression level (row-wise Z score of log2 (TP10K + 1); color scale) of genes (rows) across cells (columns) is shown sorted by subset. (C) Dot plot showing log-transformed average expression levels (dot color) and -cell percentage (dot size) for selected epithelial and endothelial genes (columns) in the different clusters (rows), for *n*=4 adult mouse lung samples. Cluster 5 (highlighted with blue frame) expresses epithelial and endothelial canonical marker genes. (D, E) Subsets of CD326+CD31+ cluster 5 cells (586 cells). A UMAP visualization (D) and dot plot showing the fractions of expressing cells (dot size) and mean expression level in expressing cells (dot color) of selected marker genes (columns) across the three subsets (rows) (E). (F-H) Pseudotime analysis of cluster 5 cells. (F) Differentiation trajectory of cluster 5 subsets. The black line is the inferred differentiation trajectory (Methods). (G) Pseudotime score depicted on UMAP visualization. (H) Pseudotime score (x-axis) of the subsets (y-axis) are distributed (with mean±SD for subclusters).

### Fate tracing reveals differentiation of lung cells from VEcad-GFP donors towards epithelial fate

To further interrogate the potential multi-potency of the patch forming cells, we used VEcad-Cre mTmG and VEcad-Cre nTnG transgenic mice expressing GFP under the VE-cadherin (Cadherin 5) promoter. Transplantation of unsorted lung cells from such GFP+ donors led to green patches in which GFP+ endothelial and epithelial cells could be clearly identified (Fig. 8), further supporting the possibility that the patch forming lung cell is expressing a canonical endothelial marker such as VE-cad, and its ability to generate epithelial AQP-5 + AT1 and Lamp-3+ AT2 alveolar cells.

**Fig. 8.**
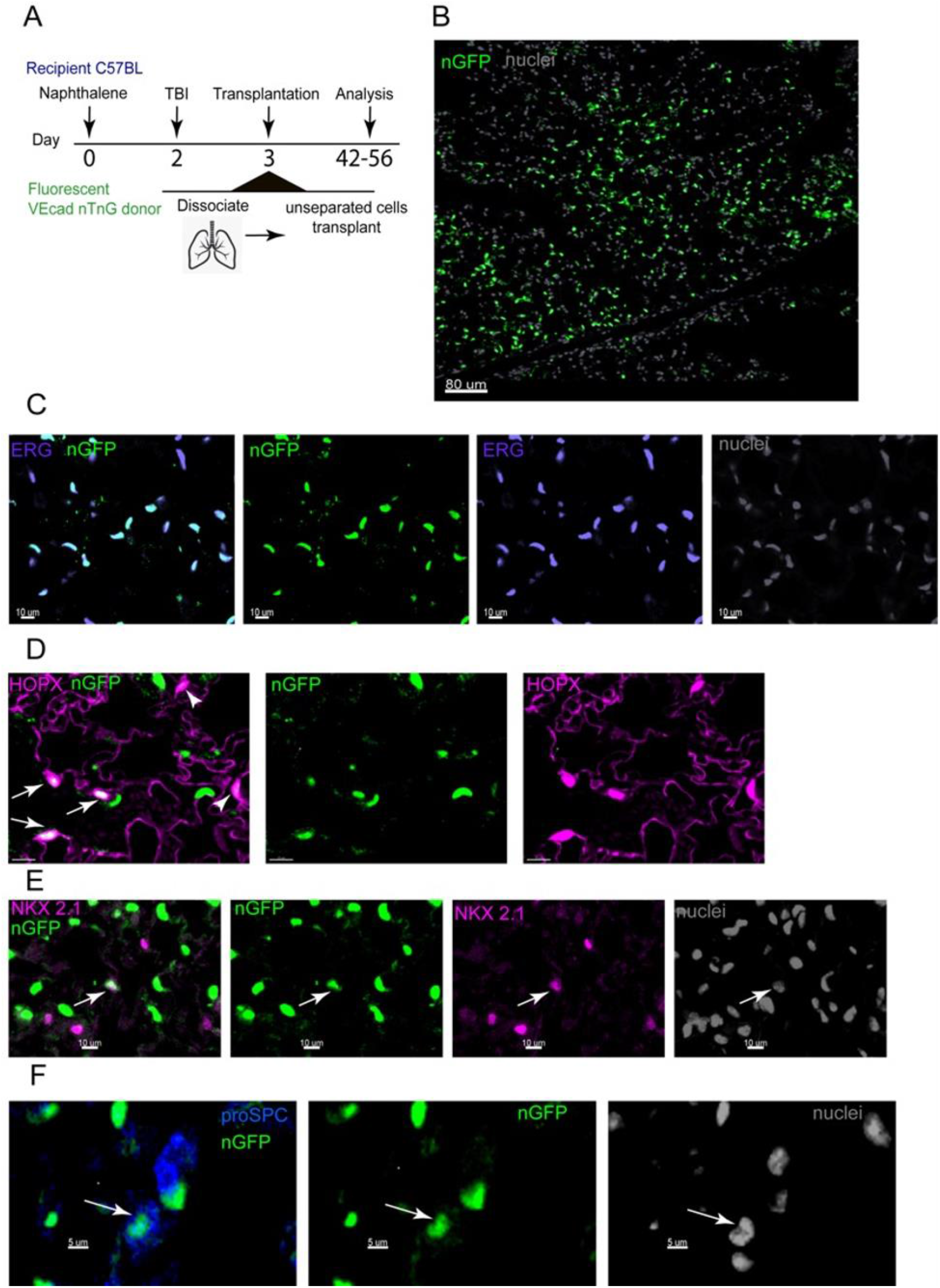
Staining of nGFP positive lung patches derived from transplanted VEcad Cre nTnG lung cells for endothelial and epithelial lineage differentiation markers. (A) Experimental design of the representative experiment. (B) Image shows nGFP-positive engrafted areas of the lung; scale bar=80 um. (C) Staining of the chimeric lung for ERG (violet), nGFP (green) and nuclei (grey); scale bar=10 um. (D) Close-up of nGFP+ AT1 donor-derived cells stained for HOPX (magenta); scale bar=5 um. Arrows indicate donor-derived nGFP+ AT1 cells and arrowheads indicate recipient nGFP-AT1 cells. (E) Donor-derived nGFP+ cell stained for Nkx2.1 (magenta); scale bar=10 um. Arrow indicates nGFP+ Nkx2.1+ cell; scale bar=10 um. (F) Donor-derived nGFP+ AT2 (arrow) cell stained for pro-SPC (blue); scale bar=5 um. The images are representative of *n*=3 mice.

### Identification by immunohistology of double positive lung cells in the mouse lung expressing canonical epithelial and endothelial markers

As CD326 staining of frozen or fixed lung tissue is not effective and based on our scRNA-seq data, suggesting that canonical epithelial gene markers such as E-Cad and NKx2.1 can be used in combination with the endothelial marker VECad to distinguish the double positive cells from single positive epithelial or endothelial cells, we analyzed lungs of transgenic mice expressing GFP in the nucleus under the VECad promoter for the presence of putative double positive progenitors. Thus, as shown in Fig. 9A, VECad (nGFP+) Nkx2.1+SOX17+ cells could be clearly identified, predominantly in proximity to blood vessels. The expression of epithelial and endothelial markers could also be ascertained by clear identification of VECad (nGFP+) Nkx2.1+CD31+ cells although nuclear staining for Sox17 is generally more reliable. For identification of these triple positive cells, we performed high-resolution confocal microscopy and traced single VECad (nGFP+) NKx2.1+SOX17+ cells along the z-axis, and verified that indeed the VE-Cad (nGFP), NKx2.1 and SOX17 markers are expressed in the same cell (Fig. 9B and Supplementary Fig.10). Enumerating these triple-positive cells in 10 fields revealed a frequency of about 0.04 % of these triple positive cells out of all counted lung cells. This low frequency aligns with the incidence of cluster 5 cells found by the scRNA-seq analysis of FACS-sorted cells. Considering that the average frequency of CD326+CD31+ cells is 3 % out of the CD45- cells (average 35% of average 91% single cells), we estimate that the frequency of the double positive cells of cluster 5 (9.2% of the sorted CD326+CD31+ population after doublets removal) in the entire lung is around 0.08%. This low incidence is similar to that that found for the triple positive VECad (nGFP+) NKx2.1+CD31+ cells in the entire lung by immunohistology. Thus, both assays reveal the presence in the lung of a unique cell expressing both epithelial and endothelial canonical genes, confirming that the identification by FACS analysis of lung cells with dual expression of epithelial and endothelial genes is not merely an artifact. Further sorting and transplantation studies using lung cells of different transgenic mice, to further characterize the patch forming subpopulation, are warranted.

**Fig. 9.**
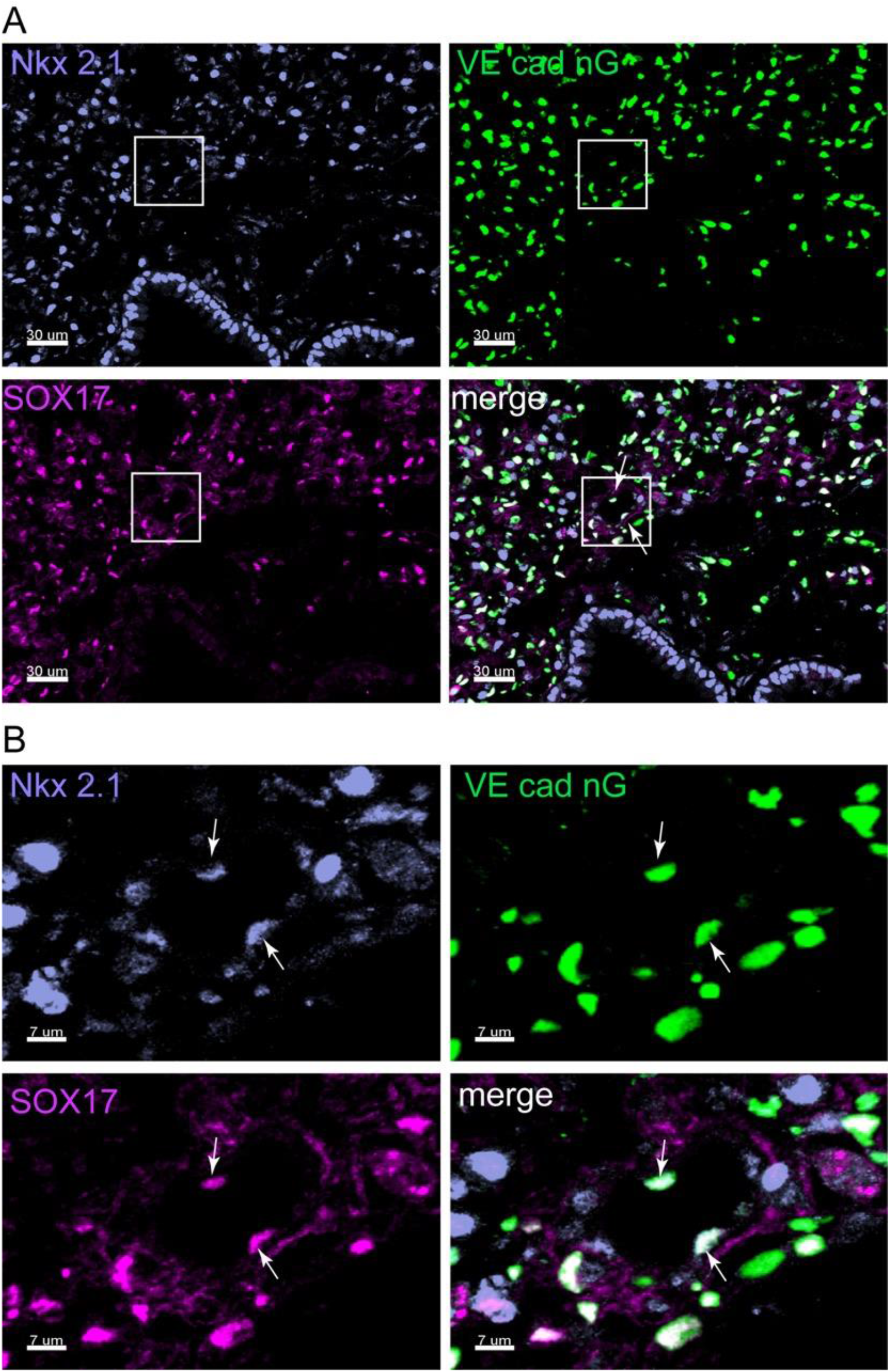
Typical Immunohistochemistry of a lung from transgenic VEcad nTnG mouse. (A) VEcad expression is depicted by nGFP (green). Expression of Nkx2.1 and SOX17 is shown with anti-Nkx2.1 (violet), and anti-SOX17 (magenta) antibody. Triple positive cells are indicated by arrows. Scale bar=30 um. (B) A close-up of the boxed area in (A). Scale bar=7 um.

## Discussion

We recently demonstrated that 6-8 weeks after transplantation of mouse or human fetal lung cells, harvested at the canalicular stage of gestation, into mice preconditioned with Naphthalene and 6Gy TBI, numerous donor-derived patches comprising epithelial and endothelial cells could be detected throughout the lung (5). These observations were recently confirmed by Shiraishi et al., following injury with elastase and 6Gy TBI (20). Similar findings were subsequently demonstrated following transplantation of adult lung cells, though the concentration of patch-forming cells was 3-4-fold lower compared to that in the fetal lung (6). Considering that the lung patches are observed within confined borders similar to spleen colonies known to be derived from a single multi-potent hematopoietic progenitor, we hypothesized that these lung patches might represent a clonal derivation from a yet unknown multi-potent lung progenitor. Here, we show, using *R26R-Confetti* donors, in which each fluorescent cell is labeled in one of four potential colors, by virtue of Cre-recombination, that at eight weeks after transplantation, all observed patches are monochromatic. This finding indicates that each donor-derived lung patch found after transplantation of fetal or adult mouse lung cells is formed by colonization and differentiation of a single multi-potent lung progenitor.

Furthermore, this finding enabled us to characterize the putative multi-potent lung progenitor by FACS purification. Thus, using our transplantation assay, we found two distinct patch-forming progenitors within the non-hematopoietic CD45- lung compartment, namely, CD326+ CD31+ and CD326-CD31+ cells. However, the single-positive CD31+ cells are predominantly associated with forming endothelial cells, while the double positive population is associated with numerous patches comprising both epithelial and endothelial cells.

While our sorting experiments strongly suggest that the multipotent patch-forming lung progenitors are found within the CD45-CD326+CD31+ cell population, our FACS data also demonstrate marked heterogeneity of CD326 expression in this lung cell subpopulation. Notably, scRNA-seq analysis of sorted CD45-CD326+CD31+ cells revealed a range of both epithelial and endothelial subsets, among them a unique cluster 5, exhibiting the expression of canonical epithelial and endothelial genes. This subset was the only one identified in our single cell analysis of either single-positive (CD326+CD31- or CD326-CD31+) or double positive (CD326+CD31+) cells to express both endothelial and epithelial markers. Pseudotime analysis showed the continuum of cluster 5 from one subset with low expression of both epithelial and endothelial markers to two subsets with higher dedication toward a specific lineage. Furthermore, doublet analysis identified only 0.3% (2 cells out of 586 cells) doublets in cluster 5. Thus, our single cell analysis suggests the existence of a bi-directional progenitor in the CD326+CD31+ population (about 9.2 % of the cells). Thus, based on our sorting strategy the cells in cluster 5 represent about 0.08% of the entire lung cell population. A similar low percentage of lung cells expressing VEcad, Sox17 and NKx2.1 could also be detected by immunohistology of the entire mouse lung of transgenic Vecad-Cre-nTnG mice expressing GFP under the VE-cadherin (Cadherin 5) promoter.

Notably, as indicated by immunohistology, these cells are concentrated around blood vessels similar to hematopoietic progenitors residing in a vascular niche in the bone marrow (21). Further analysis of cluster 5 identified a continuum of cellular heterogenicity with three subsets expressing canonical epithelial and endothelial gene markers along the trajectory of differentiation, bifurcate to endothelial or epithelial lineages. Thus, examining the endothelial and epithelial double-positive lung population by scRNA-seq further supports our *in vivo* results by identifying a small subset of cells that might account for early bidirectional lung progenitor.

Additional support for this hypothesis was found by fate tracing experiments using transplantation of lung cells from VEcad-Cre-mTmG and VEcad-Cre-nTnG transgenic donors. Thus, these transplants led to green lung patches comprising epithelial and endothelial cells expressing GFP, likely originating from a lung progenitor expressing a canonical endothelial marker such as VEcad, and capable of differentiating along both epithelial and endothelial lineages. It could be argued that prior to transplantation, GFP staining might leak in the VEcad-GFP donor mice into a small number of epithelial CD326+CD31-cells. However, considering that our sorting experiments rule out a patch-forming activity in the single positive CD45-CD326+CD31-cells or the ability to generate CD326+ progeny from the single positive CD45-CD326-CD31+ cells, the results of these transplantation experiments are most consistent with the presence of a multi-potent lung progenitor within the double positive CD326+CD31+ population.

A classic example of such cellular plasticity was demonstrated by Tata et al. (22, 23), showing de-differentiation of fully mature secretory cells into basal stem cells with regenerative capacity. Another example of cellular plasticity, known as trans-differentiation or trans-determination, occurs when stem cells from one region of the lung can convert into stem cells associated with the other lung region (23). Moreover, evidence of multi-potential progenitors capable of developing into endothelial and epithelial lineages, has also been described for human breast progenitors, which were found to express a significant level of ‘Yamanaka’ transcription factors (24, 25).

Collectively, all three assays used to characterize the cell fraction leading to multipotent patches upon transplantation, support the presence in this heterogenic lung cell fraction of a unique cell expressing canonical genes characteristic of both epithelial and endothelial lineages. It could be argued that the sorted CD45-CD326+CD31+ cell fraction might represent CD326+CD31- epithelial cells that somehow attach to CD326+CD31- cells and form cell doublets, and that upon transplantation of such doublets the formed donor-derived patches include epithelial cells originating from CD326+ CD31- progenitors and the endothelial cells originate from CD326-CD31+ progenitors. However, this potential interpretation is negated by three major findings: 1) The patches derived from transplantation of cells from *R26R-Confetti* donors lead to monochromatic patches. 2) Transplantation of sorted Td-Tomato CD326+CD31-cells did not lead to any donor-derived Td-Tomato patches in our assay, even if combined with green unseparated lung cells (including CD326-CD31+), while this combined cell fraction leads to the formation of green patches. 3) Single cell analysis of double positive cells but not in single positive cells, identify a small subset of cells (about 9% of the double positive cells, cluster 5) that express both epithelial and endothelial markers, strengthening our transplantation results for the existence of an early bi-directional progenitor cell within the double positive heterogenous population. Doublet finder analysis of the scRNA-seq data showed very low frequency of doublets in the double positive cluster 5.

Thus, further studies using transgenic mice to identify additional markers of the putative CD45-CD326+CD31+ lung progenitor are warranted. For example, it is possible to generate a genetic lineage-tracing system using dual recombinases (Cre-FLP) (26), to specifically track and sort the putative patch-forming lung progenitor population using markers identified by scRNA-seq in cluster 5 (e.g., Cdh5 and Nkx2.1).

In summary, based on the use of *R26R-Confetti* donors, we have established that donor-derived patches found after transplantation of lung cells into lung-injured recipient mice, originate clonally from a single progenitor. In addition, FACS sorting data show that these progenitors are included within the CD45-CD326+ CD31+ lung cell sub-population, which is distinct from CD45-CD326-CD31-, CD45-CD326-CD31+ or CD45-CD326+CD31-cells, in its capacity to form patches comprising both epithelial and endothelial cells. This multi-potent capacity of the patch-forming cells is in striking contrast to previously described lung progenitors that are restricted to differentiation along epithelial lineages (1), (2). Such pluripotency could be of particular value for lung injury repair, considering that all the major lung diseases exhibit not only epithelial injuries, but are also associated with endothelial damage.

Furthermore, apart from its potential for translational studies aiming at the correction of different lung diseases, the identification of novel lung progenitors could also contribute to basic studies aimed at a better understanding of fetal lung development, as well as an understanding of steady-state maintenance of different cellular lineages in the adult lung.

## Materials and Methods

### Mice

Animals were maintained under conditions approved by the Institutional Animal Care and Use Committee at the Weizmann Institute and MD Anderson Cancer Center. All of the procedures were monitored by the Veterinary Resources Unit of the Weizmann Institute and MDA and approved by the Institutional Animal Care and Use Committee (IACUC). Mice strains used included: *Rag1*^-/-^ (on C57BL/BJ background), C57BL/6J (CD45.2) and C57BL/6-Tg (CAG-EGFP)1Osb/J, (Weizmann Institute Animal Breeding Center, Rehovot, Israel), Gt(ROSA)26Sortm4(ACTB-tdTomato,-EGFP)Luo/J (27), B6.129P2-Gt(ROSA)26Sortm1(CAG-Brainbow2.1)Cle/J (9), B6.129-Tg(Cdh5-cre)1Spe/J (28), B6N.129S6-*Gt(ROSA)26Sor^tm1(CAG-tdTomato*,-EGFP*)Ees^*/J (29), Nkx2-1tm1.1(cre/ERT2)Zjh/J (30), (Jackson Labs, Bar Harbor, USA). All mice were used at 6–12 weeks of age. Mice were kept in small cages (up to five animals in each cage) and fed sterile food and acid water. Randomization: Animals of the same age, sex and genetic background were randomly assigned to treatment groups. Pre-established exclusion criteria were based on IACUC guidelines, and included: systemic disease, toxicity, respiratory distress, interference with eating and drinking, substantial (above 15%) weight loss. During the entire study period, most of the animals appeared to be in good health and were included in the appropriate analysis. In all experiments, the animals were randomly assigned to the treatment groups.

### Naphthalene lung injury

Naphthalene (> 99% pure; Sigma-Aldrich) was dissolved in corn oil, and administered at a dose of 200 mg per kg body weight, by intra-peritoneal injection, 40–48 hrs before exposure to TBI, as previously described (5–7) For “double” lung injury, naphthalene treated animals were further irradiated in an X-Ray Irradiator (40–48 hrs after naphthalene administration). C57BL/6, and *Rag1*^−/−^ mice were irradiated with 6Gy TBI.

### Fetal and adult lung cell transplantation

Fetal and adult lung cell suspensions were obtained by enzymatic digestion, as previously described (5–7), with some modifications. Briefly, lung digestion was performed by either finely mincing tissue with a razor blade in the presence of 1% collagenase, 2.4 U/ml Dispase II (Roche Diagnostics, Indianapolis, IN), and 1 mg/ml DNAse-I (Roche Diagnostics, Indianapolis, IN) in PBS Ca^+^Mg^+^, or by enzymatic digestion of the lung tissues in the presence of 1mg/ml collagenase, 2.4 U/ml Dispase II and 1 mg/ml DNAse-I (Roche Diagnostics) in Ca^+^Mg^+^ phosphate buffered saline in GentleMACS™ Octo Dissociator with Heaters (Miltenyi Biotec), using the mouse lung dissociation protocol provided by the vendor. Removal of nonspecific debris was accomplished by sequential filtration through 70- and 40- μm filters. The cells were then washed with PBS including 2% FCS, 2mM EDTA, and antibiotics.

C57BL/6 mice or Rag1^-/-^ recipient mice were conditioned with naphthalene, and after 48 hrs, exposed to 6Gy TBI. The mice were transplanted with 1x10^6^ E15– E16 embryonic or 3-16x10^6^ adult mouse lung cells by injection into the tail vein 4–8 hrs following irradiation.

### TMX administration

TMX (Sigma-Aldrich) was prepared in corn oil, as a 20mg/ml stock solution. For induction of Cre recombination in *R26R-Confetti* adult mice, the mice were injected intraperitoneally (IP) with two 5mg doses of TMX, - on days 5 and 4 prior to harvest of bone marrow or lungs. Lungs or BM were harvested 5 days after TMX administration. For induction of CRE recombination in Confetti embryos, female mice were treated at E12 with single IP dose of 5mg TMX. The embryos were harvested at E16, and lung and liver cells of the Confetti-positive embryos were used for transplantation experiments.

### Flow cytometry analysis of *R26R-Confetti* donor cells and transgenic mouse lungs

FACS analysis was performed on LSRII (BD Biosciences) or Fortessa analyzers equipped with 5 lasers. E16 fetal Confetti lung and liver cells, and adult Confetti lungs and bone marrow cells were analyzed. E16 fetal and adult lung cells were analyzed for YFP/GFP, RFP and CFP labeled cells as well as for epithelial (CD326), endothelial (CD31), and hemopoietic (CD45) lineage markers, to quantify the different cell populations within the cells marked with different fluorescent tags after Cre recombination, induced by TMX administration.

Samples were stained with the conjugated antibodies or matching isotype controls according to the manufacturer’s instructions. The full list of anti-CD326, anti-CD31 and anti-CD45 antibodies is provided in Supplementary Table 5. E16 fetal liver and adult bone marrow cells were analyzed for the Lin^-^ Sca-1^+^c-kit^+^ cell (LSK) population. The cells were stained with the following antibodies or matching isotype controls: Lineage panel-streptavidin, followed by staining with biotin -APC or biotin APC-CY7, Sca-1-Brilliant Violet 711, and C-kit PE-CY7.

A full list of antibodies used is provided in Suppl. Table S1. Antibodies were purchased from e-Bioscience, BD, and Biolegend.

Data were analyzed using FlowJo software (version vX.0.7 Tree Star Inc)

### Image acquisition by two-photon laser scanning microscope (TPLSM)

A Zeiss LSM 880 upright microscope fitted with Coherent Chameleon Vision laser was used for imaging experiments of explant lung tissue. Images were acquired with a femtosecond-pulsed two-photon laser tuned to 940 nm. The microscope was fitted with a filter cube containing 565 LPXR to split the emission to a PMT detector (with a 579-631 nm filter for tdTomato fluorescence) and to an additional 505 LPXR mirror to further split the emission to two GaAsp detectors (with a 500-550nm filter for GFP fluorescence). Images were acquired as 100 -150 µm Z-stacks with 1-5 µm steps between each Z-plane. The zoom was set to 0.7, and pictures were acquired at 512 x 512 x-y resolution.

### Tissue clearing

Mice were perfused with monomer solution containing 4% PFA, 4% acrylamide, 0.0125% bis-acrylamide, and 0.1% azo-initiator, VA-044. The lungs were immediately immersed in the above solution for an additional 24-48h at 40C with constant shaking, to avoid premature polymerization. After degassing, the lungs were left to polymerize for 3h at 37°C and washed with 20mM sodium borate buffer containing 200mM SDS at pH=9. The lungs were then cleared for 4 days using a rotational electrophoresis clearing system (the SmartClear II Pro, Life Canvas Technologies, Seoul, South Korea). Once the lungs reached a sufficient clearing state, they were washed in 20mM sodium borate buffer for 24h and immersed in index matching solution until imaging (EasyIndex; RI=1.46, Lifecanvas technologies).

### Light-Sheet microscopy

For imaging of large lung volume, three-dimensional images of cleared lungs were acquired using an ultramicroscope II (LaVision BioTec GmbH, Astastraße 14, 33617 Bielefeld / Germany) operated by the ImspectorPro software (LaVision BioTec). The light sheet was generated by a Superk Super-continuum white light laser with an emission range of 460-800nm, mW/nm (NKT photonics, Blokken 84, DK-3460 Birkerød). Excitation filters used to detect red and green patches were 545/25 and 470/40, respectively. The corresponding emission filters were 595/40 and 525/50. The microscope was equipped with a single lens configuration, using a 4X objective (LVMI-Fluor 4X/0.3; NA: 0.3; WD: 5.6-6.0 mm; RI range: 1.33-1.57). Samples were glued to the sample holder and placed in an imaging chamber made of 100% quartz (LaVision BioTec) filled with EasyIndex solution (RI=1.46; Lifecanvas technologies) and illuminated from the side. Images were acquired by an Andor Neo sCMOS camera (16bit, 2150 × 2560, pixel size 1.626 x 1.626 µm, Andor 277.3 mi • Belfast, UK). Z stacks were acquired with 5μm steps, and two fields of view, 3510×4160 µm each, were acquired with 20% overlap, and stitched using Imaris stitcher (BITPLANE by Oxford Instrument, http://www.bitplane.com).

To increase the imaging resolution of the red and green engrafted lung patches in 3D, samples were also imaged using a LightSheet Z.1 microscope equipped with two PCO.Edge, 1920×1920 sCMOS cameras (Zeiss Ltd, Tegeluddsvägen 76 115 28 Stockholm, Sweden). The red and green patches were illuminated using Zeiss illumination optics lightsheet 10X/0.2, and their emission detected using Clr Plan-Neofluar 20×/1.0 Corr nd=1.45 (Zeiss Ltd.). Samples were glued at their edge to a holder, and immersed into the imaging chamber, filled with EasyIndex solution (Lifecanvas Technologies).

Imaging was performed using single side illumination, at two fields of view: 1093.91×1093.91×296.22 µm, and 437.56×437.56×315.33 µm. The excitation lines for the red and green patches were 561nm at 1% and 488nm at 2%, with collected emission at 575-615nm and 505-545nm, respectively.

### Assessment of *R26R-Confetti* ^+^ foci in chimeric lungs by immunohistology

Lungs were fixed with a 4% PFA solution introduced through the trachea under a constant pressure of 20 cm H_2_O. Then, the lungs were immersed in fixative overnight at 4°C. Lungs were processed after PFA treatment, fixed in 30% sucrose, and frozen in Optimal Cutting Temperature (OCT) compound (Sakura Finetek USA, Inc.Tissue-Tek). Serial step sections, 12 μm in thickness, were taken along the longitudinal axis of the lobe. The fixed distance between the sections was calculated to allow systematic sampling of at least 20 sections across the entire lung. Lung slices were analyzed by fluorescence or confocal microscopy. The actual number of “Confetti” patches (a group of more than five adjacent cells labelled with the same fluorescent tag: cytoplasmic RFP or YFP, nuclear GFP, or membrane CFP, was defined as a single patch) was counted per slice.

### Confocal Microscopy

Thin 12µm sections of engrafted lung were imaged using an upright laser scanning Leica TCS SP8 microscope, equipped with acousto optical beam splitter and acousto optical tunable filter (Leica microsystems CMS GmbH, Germany) for wavelength separation, and two internal HyD detectors equipped with spectral separation. The confocal pinhole was open to 1AU (58.6 µm for 580 nm). Images were acquired using the 8k Hz resonant scanner in a 1024*1024, 8bit format at two magnifications. Use of 20X air objective (HC PL 20x/0.75 W, Leica microsystems), provided images with a field of view of 443.29µm; pixel size=0.43 µm. For higher resolution images, a 60X oil objective was used (HC PL APO 63x/104 oil CS2, Leica microsystems) providing image dimensions of FOV=140.73 µm; pixel size= 0.137 µm.

To attain well distinguished excitation and signal collection from the five different markers (see the list of antibodies and staining details) sequential imaging steps were applied, with sequence shift following each Z stack acquisition: The 1^st^ sequence applied excitation with an Ar laser at 2% (of 30% laser) at 488nm, and HeNe633 laser at 1%, with emission collected at 505-566nm and 651-708nm, respectively. The 2^nd^ sequence applied excitation using a DPSS561 laser at 4% and collection at 574-638nm. The 3^rd^ sequence applied excitation with a Diode405 laser at 8% and collected two emissions at 413-446nm and 493-521nm.

### Wide-Field Microscopy

To image the entire area of a lung tissue section, a Leica DMi8 inverted microscope was used, equipped with a motorized stage for fast imaging. Imaging was done with a 10X air objective (HC PL FLUOTAR 10x/0.3 DRY, Leica microsystems) and recorded with a CCD camera (1392×1040, 8bit, Leica DFC7000 GT monochromatic, Leica microsystems).

### Transplantation of sorted CD326+CD31+, CD326+ CD31-, CD326-CD31+ cell populations into NA+6Gy TBI preconditioned mice

Experiments were performed involving transplantation of FACS-sorted cell populations from adult mouse lungs. The labeled donor mice used included GFP (C57BL/6-Tg (CAG-EGFP)1Osb/J), mTmG (*Gt(ROSA)26Sor^tm4(ACTB-tdTomato,-EGFP)Luo^*/J_) and nTnG *(Gt(ROSA)26Sor*_*tm1(CAG-tdTomato*,-EGFP*)Ees*_/J) (mTmG mice_ express membrane td-tomato, and nTnG mice express nuclear td-tomato). Altogether, a total of 16 independent transplantation experiments were performed, testing the patch-forming activity of sorted cells. Live, single CD45- lung cells were FACS sorted into four subpopulations including CD326+ CD31-, CD326+CD31+, CD326- CD31+ and CD326-CD31- cells. The 0.3-0.5 x10^6^ sorted cells from either mTmG or nTnG donors were transplanted with or without 0.5 x10^6^ unseparated lung cells from GFP+ mouse donors into naphthalene treated and irradiated C57BL mice. The lungs of transplanted mice were harvested and evaluated for the presence of donor-derived cells 6 to 24 weeks post-transplantation.

### Single-molecule fluorescent *in situ* hybridization (smFISH)

SmFISH was performed as previously described (31) For smFISH, lung tissues were harvested, inflated, and fixed in 4% paraformaldehyde for 3 hours; samples were incubated overnight with 30% sucrose in 4% paraformaldehyde, and then embedded in OCT and frozen. 6 µm cryosections were used for hybridization. smFish probes were coupled to Cy5. The probes were purchased from Stellaris® Biosearch^TM^ Technologies.

### Single-cell RNA-Seq analysis of FACS purified CD326+CD31+, CD326+CD31- and CD326-CD31+ cells

Single cell RNA transcriptome analyses were performed on CD326+CD31+, CD326+CD31-, and CD326-CD31+ datasets. The lungs from WT mice were enzymatically treated, dissociated into single cells, pooled, and FACS sorted for CD45-CD326+CD31+, CD45-CD326+CD31-, and CD45-CD326-CD31+ cells. The single cell RNA transcriptome analysis was performed at the MD Anderson core lab, using the 10x Genomics platform. The Cell Ranger Single Cell Software Suite v3.0.1 (https://support.10xgenomics.com/single-cell-gene-expression/software/overview/welcome) was used for sample demultiplexing, alignment to the mm10 mouse reference genome and transcriptome, and for generating filtered unique molecular identifier (UMI) count matrices, which were used for downstream analyses. The R package, Seurat v3.1.1 (32–34) was used to perform data QC, normalization, and integration of the three datasets. Cells with dataset-specific outliers of high mitochondria percentage, extremely high or low number of genes, or high RNA content were filtered out as potential dead cells or doublets. LogNormalize was used for normalization. Integration was based on 2000 anchor genes as a default. The R package monocle3 v0.2.0 (35–37) was used for dimension reduction of the integrated dataset using the uniform manifold approximation and projection (UMAP) method (38) and unsupervised clustering of cells was performed using Louvain/Leiden community detection (39). Marker genes of each cluster were identified using Seurat by comparing the gene expression profile of each cluster to all others using Wilcoxon rank sum test. A single-sample gene set variation analysis was performed to calculate cluster-wise gene set enrichment scores using the R package, GSVA (40) v1.32.0 on MSigDB (41) hallmark gene sets (v7.0) and curated lung gene sets (https://research.cchmc.org/pbge/lunggens/mainportal.html). Clustering of each dataset was carried out using Seurat v4.3.0. Clustree v0.5.0 (42) was used for choosing the optimal resolution. Feature and dot plots were generated to identify clusters expressing canonical epithelial and endothelial markers. DoubletFinder v2.0.3 (43) was used to identify doublet cells in the datasets. The expected doublet rates were based on number of recovered cells and 6.1%, 7.6%, and 6.9% were used for datasets CD326+CD31+, CD326+CD31- and CD326-CD31+, respectively. Heatmap of CD326+CD31+ cell-subset signatures shows the top 5 marker genes for each cluster, with cluster 5 showing an additional 5 differentially expressed genes compared to cluster 4. Marker genes for each cluster and differentially expressed genes between cluster 5 and cluster 4 were identified using Seurat v4.3.0 by Wilcoxon Rank Sum test. Clusters 1 and 2 in the heatmap were downsampled to 200 cells in order to show the smaller clusters. For further analysis of cluster 5 in CD326+CD31+ dataset, cells of this cluster were extracted and clustered in Seurat v4.3.0 and then converted to the celldataset object used by Monocle3 v1.0.0 for pseudotime trajectory analysis. Trajectory graph was learned automatically by Monocle3 and pseudotime was calculated after assigning cluster 1 as root, using default parameters.

### Statistical analysis and reproducibility

Sample size calculations were not performed. Both female and male mice were used as donors and recipients. Fetal lungs used for transplantation experiments were harvested at embryonic day E16. Adult lung donors used for transplantation of either unseparated or FACS purified populations were 6-12 weeks old. All the experiments were conducted in at least 3 biological replicates. All data are presented as the mean ± SEM or SD. Statistical analyses were performed using Prizm software. Comparisons were tested using Student’s t test, χ2 distribution test or one-way analysis of variance (ANOVA). P value of < 0.05 was considered to be significant. All graphs were generated using Excel, Prizm and Adobe Illustrator CC 2018 software.

### Supplementary Material includes

Supplementary Figures 1 to 10

Supplementary Table 1. Example of patch area and volume calculation

Supplementary Tables 2-4 (uploaded separately as exel files).

Supplementary Table 5. List of Antibodies Captions for Supplementary Movies S1 to S3

## Acknowledgments

We thank Dr. Beata Toth and Dr. Shalev Itzkovitz for assistance with setting up the Confetti system and for helpful discussions, and Dr. Shelley Schwarzbaum for critical reading of the manuscript. The imaging platform used for this work was supported by the De-Picciotto-Lesser Cell Observatory and the Elsie and Marvin Dekelboum Family Foundation.

## Author contributions

CR designed, performed and organized most of the experiments; analysed and interpreted the data; and co-wrote the manuscript.

ES analysed and interpreted the data and participated in discussions.

IMK, RO, XS, RY, MM assisted in performing experiments and analysed the data.

MS, AB, ZS, SEF participated in microscopy data acquisition, analysis and discussions.

YQ, JW assisted with computational analysis of single cell RNA seq data

MB participated in single cell RNA sequencing data analysis and critical reading of the manuscript.

YR designed, coordinated and conducted the study, including analysis and interpretation of data, and co-wrote the manuscript.

