## Supplementary Materials for "Identification of a multipotent lung progenitor for lung regeneration"

### Identification of a multi-potent lung progenitor for lung regeneration

**This PDF file includes:**

Supplementary Figures 1 to 10

Supplementary Table 1. Example of patch area and volume calculation

Supplementary Tables 2-4 (uploaded separately as exel files).

Supplementary Table 2 (Supplementary Table 2 TOP 100 genes NC Double pos)

Supplementary Table 3 (Supplementary Table 3 TOP 100 genes NC Epcam)

Supplementary Table 4 (Supplementary Table 4 TOP 100 genes NC Pecam1)

Supplementary Table 5. List of Antibodies

Captions for Supplementary Movies S1 to S3

**Supplementary Figures**

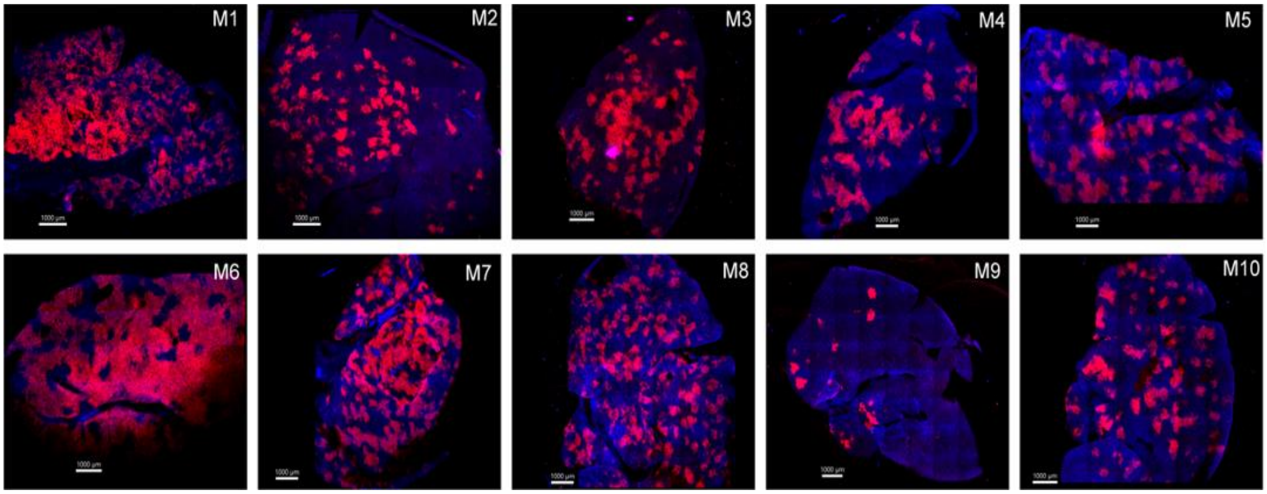

**Supplementary Fig. 1.** Chimeric lungs of mice preconditioned with NA+6Gy TBI and transplanted with TdTomato donor cells. Whole mount confocal lung images of n=10 individual chimeric mice, 8 months posttransplant.

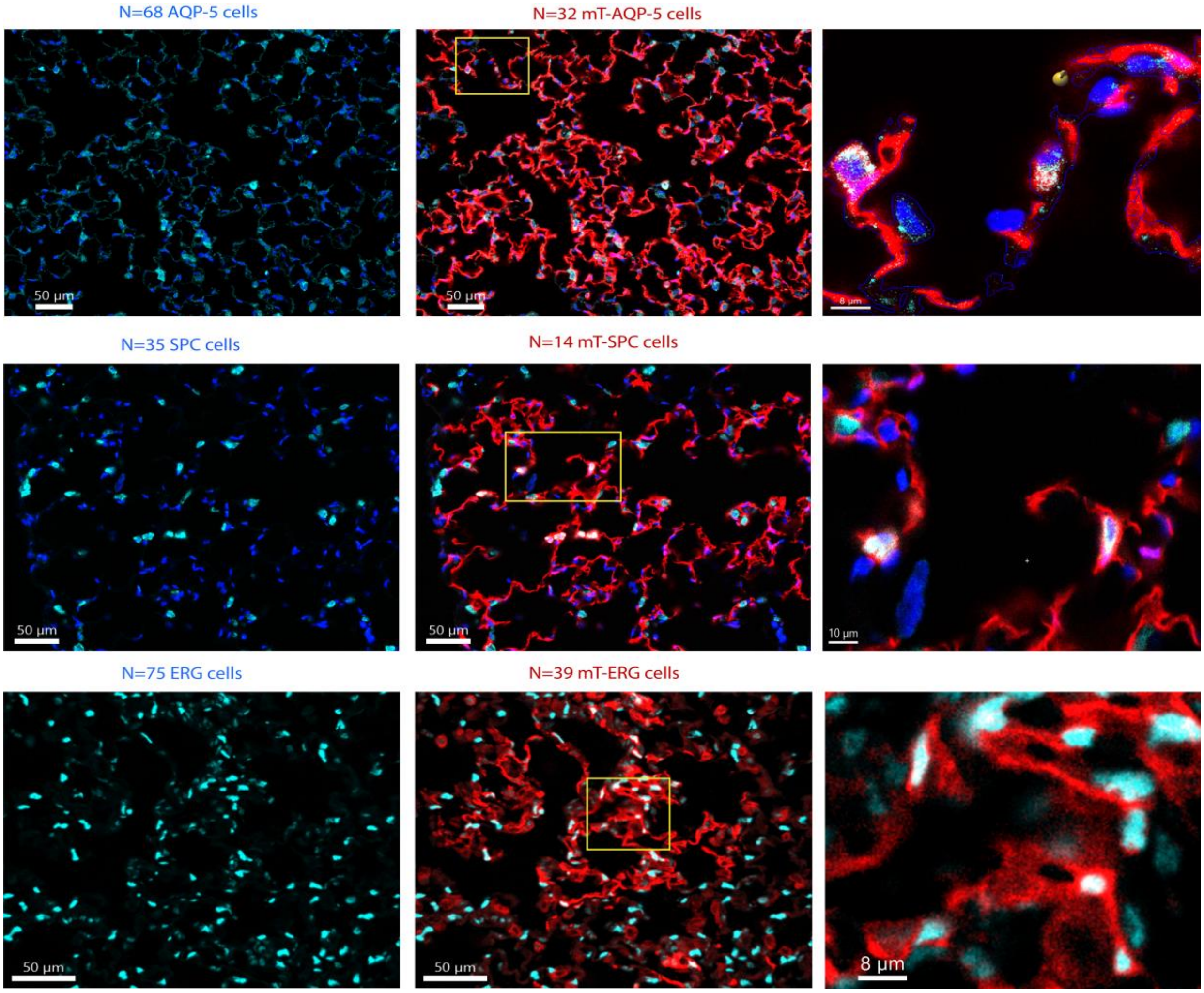

**Supplementary Fig. 2**. Images used for calculation of cell numbers of in the chimeric regions: AQP-5 positive AT1 cells, stained with smRNA probe, SPC positive AT2 cells stained with smRNA probe, ERG positive endothelial cells, stained with anti- ERG antibody (nuclear staining). Images are representative for n=2 chimeric mice. Numbers of counted cells are shown per 0.2 $mm^2$ (width of counted images were W=500um, height H= 400um respectively). Total number of AQP-5, SPC and ERG positive cells were manually counted, and then TdTomato positive cells were identified.

The percentages of donor derived AQP-5, SPC and ERG cells are 47%, 40% and 52% respectively. The right image of each row is a close up of boxed area in the middle image.

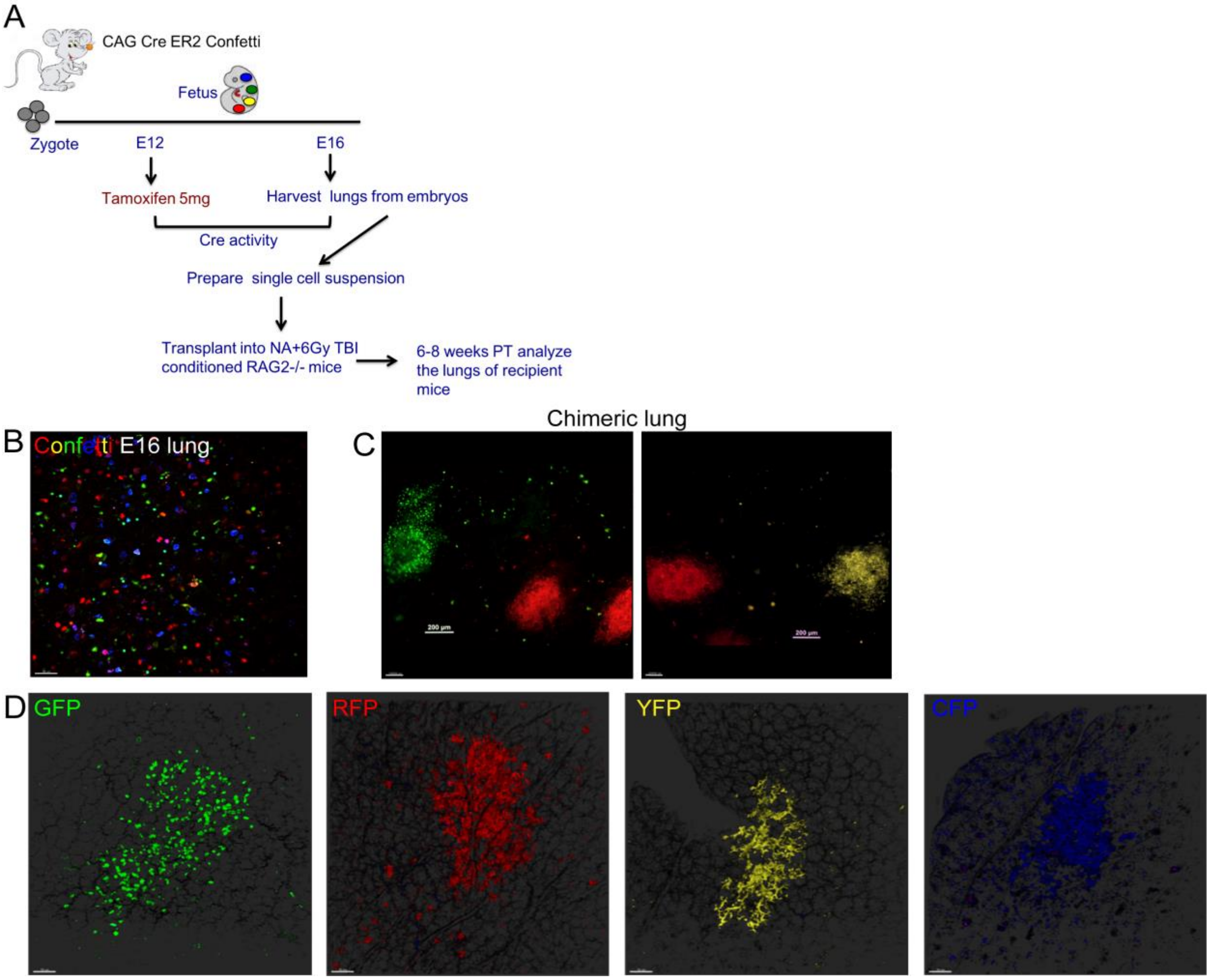

**Supplementary Fig. 3. Transplantation of fetal lung cells from *R26R-Confetti* donors.** (A) Experimental scheme using E16 fetal lung cells after induction of Cre recombination by Tamoxifen for implantation into immune deficient RAG recipients preconditioned with naphthalene and 6Gy TBI. (B) Two photon microscopy image of E16 fetal lung tissue, illustrating single monochromatic cells in *R26R-Confetti* mouse after TMX induction and prior to transplantation (from n=3 embryos). (C) Monochromatic patches at 6 weeks after transplantation of E16 fetal lung cells from *R26R-Confetti* donors into RAG2-/- recipient mouse. Whole mount of the chimeric lung was analyzed by fluorescent microscopy under low magnification (scale bar=200um. (D) Typical analysis of the same chimeric lung by two photon microscopy showing monochromatic patches expressing each of the four fluorescence tags (Scale bar=50 um). Control of non-fluorescence in other channels for RFP and CRP was negative. For GFP nuclear expression was ascertained as opposed to cytoplasmatic expression for YFP. Typical example for such a control experiment under the different channels is shown for transplantation of adult lung cells in Extended Fig.7. The results are representative of three independent experiments: n=3 mice in each experiment.

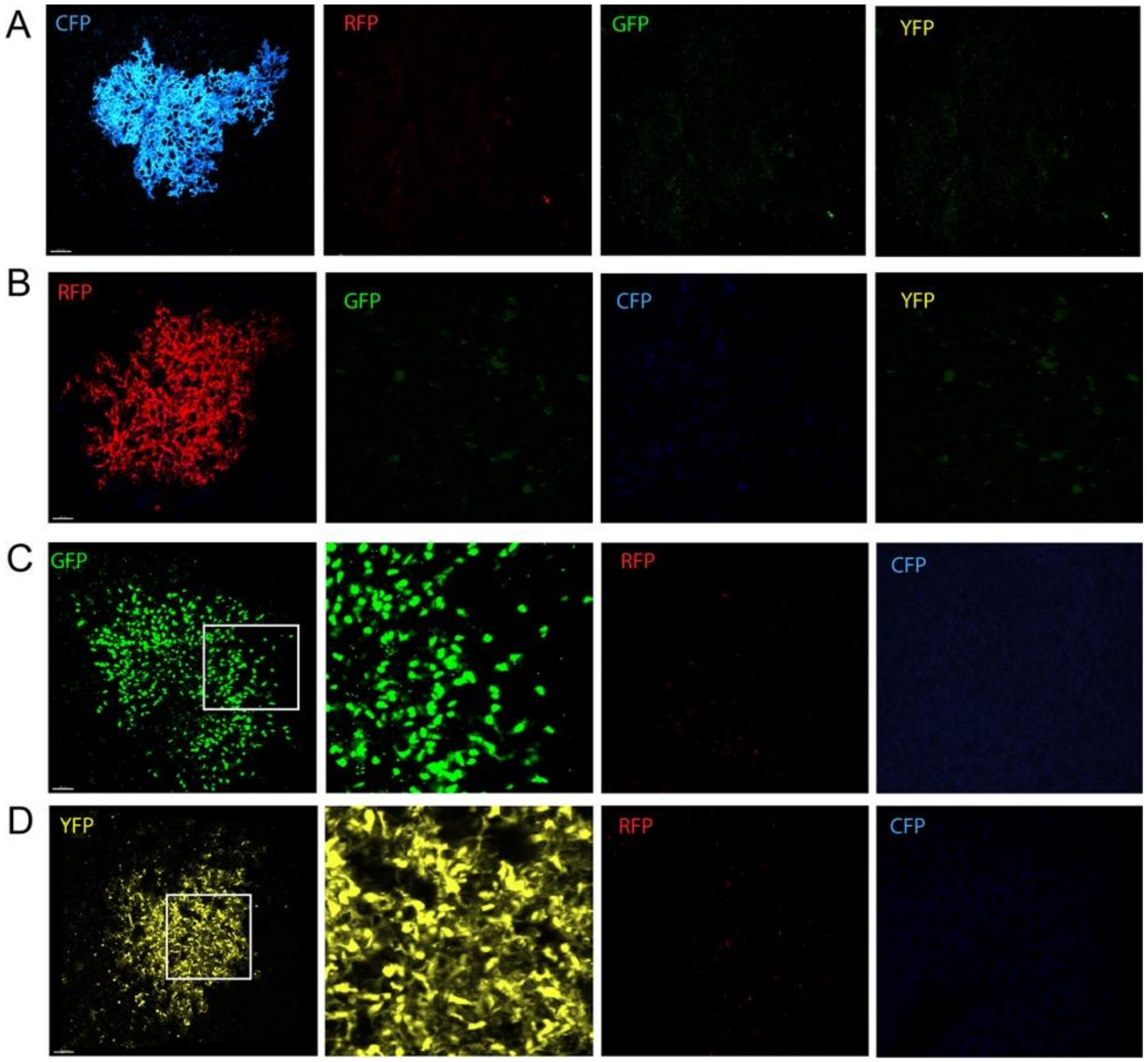

**Supplementary Fig. 4. Typical monochromatic patches, each exhibiting a distinct color in whole mount chimeric lung evaluated by two-photon microscopy.**

(A) Shows membranous CFP patch with robust fluorescence in CFP channel and complete lack fluorescence in RFP, GFP and YFP channel. (B) Shows cytoplasmatic RFP patch with robust fluorescence in RFP channel and complete lack fluorescence in CFP, GFP and YFP channel. (C) Shows nuclear GFP patch with robust fluorescence in GFP channel and complete lack of fluorescence in CFP and RFP channel. (D) Shows cytoplasmatic YFP patch with robust fluorescence in YFP channel and complete lack of fluorescence in CFP and RFP channel. YFP and GFP patches are easily distinguished by the location of the fluorescent protein within the cells- GFP in the nuclei and YFP in the cytoplasm (left middle images in C and D are zoomed in on boxed areas of the left images in C and D respectively) .

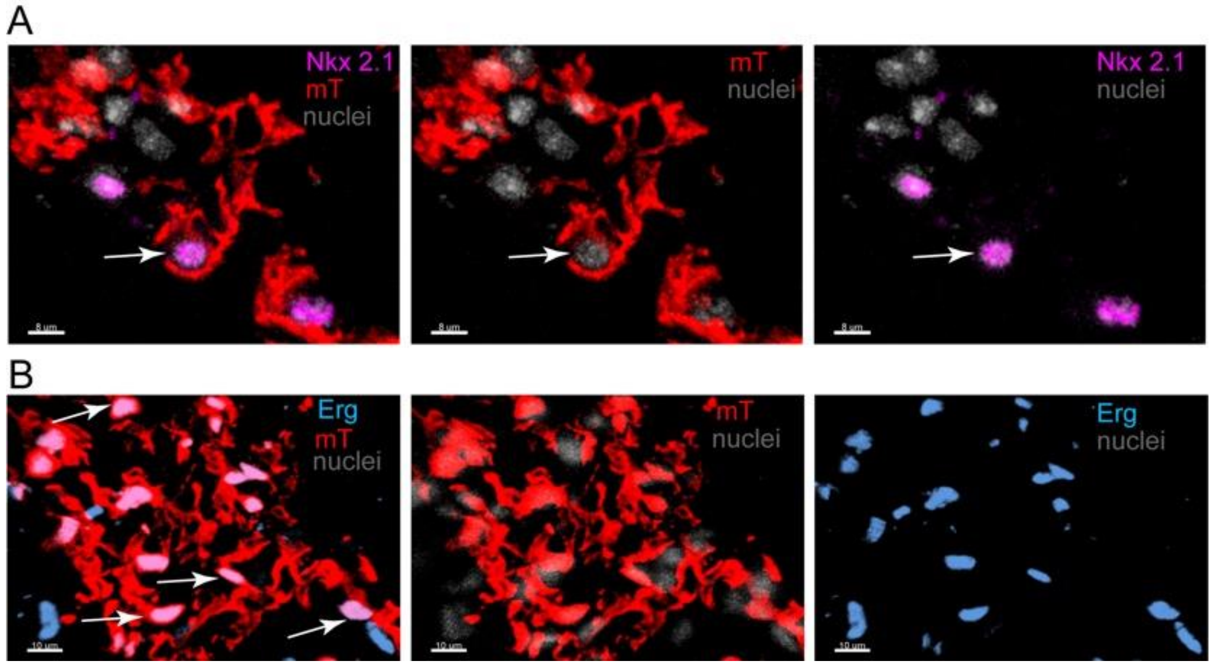

**Supplementary Fig. 5. Staining of chimeric lung, transplanted with Confetti lung cells 3 months posttransplant.** Confocal images demonstrate RFP patch stained for nuclear marker Nkx-2.1 (magenta) in (A), scale bar=8um and in (B) staining for endothelial nuclear marker Erg (cyan), scale bar=10um. The images are representative for n=2 mice.

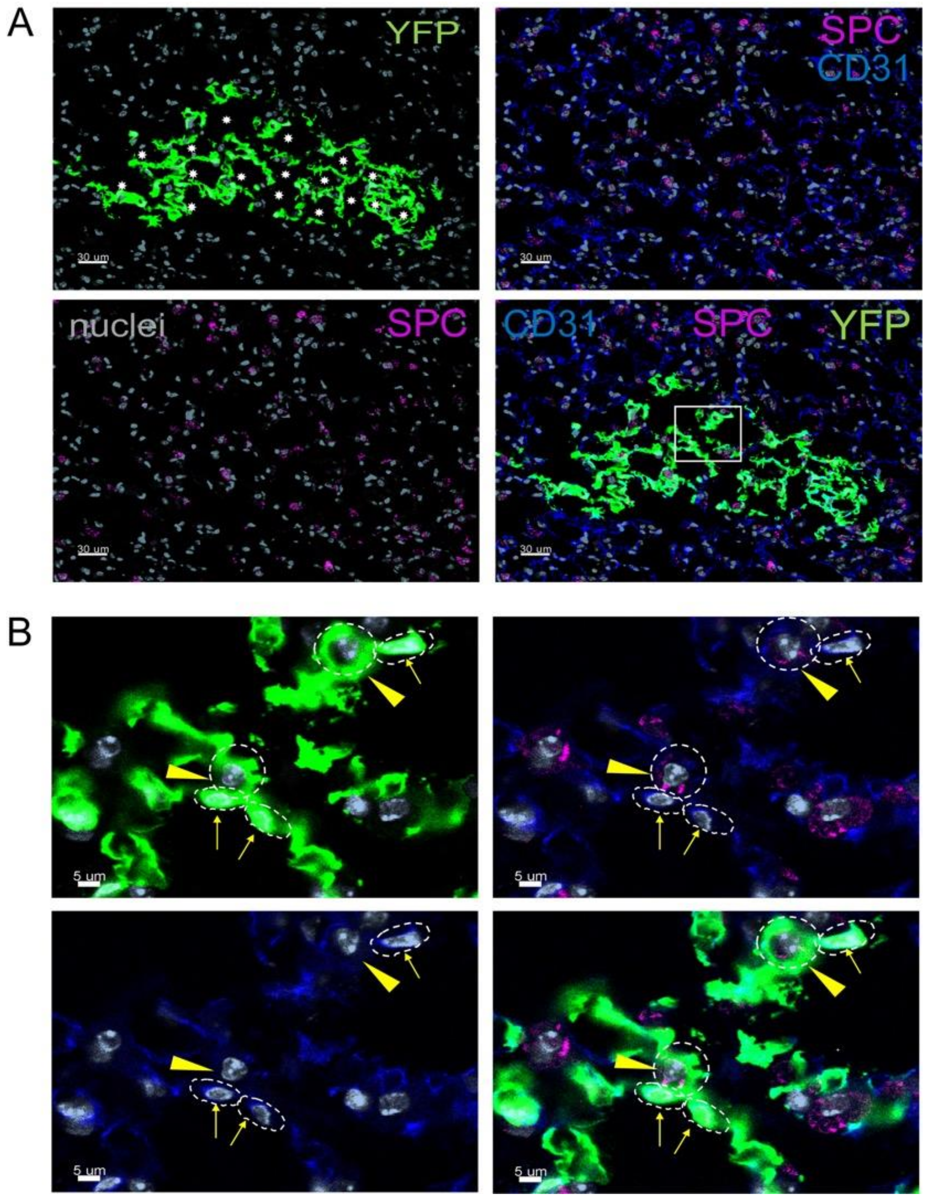

**Supplementary Fig. 6. Confocal images showing SPC+ AT2 alveolar cells and CD31+ endothelial cells within a single YFP+ patch.** (A) Typical alveoli (stars) within a YFP+ patch, showing host and donor cells stained for SPC and CD31 (Scale bar=30µm).

(B) Confocal close-ups of the squared area in (A). Hoechst nuclear staining in grey, SPC in magenta, and CD31 in blue. AT2 cells are indicated by arrow heads, and endothelial cells by arrows (Scale bar=5µm). Dotted circles mark boundaries of donor-derived AT2 cells, and dotted ellipses mark the boundaries of donor-derived endothelial cells. The images are representative for n=4 mice, from two experiments.

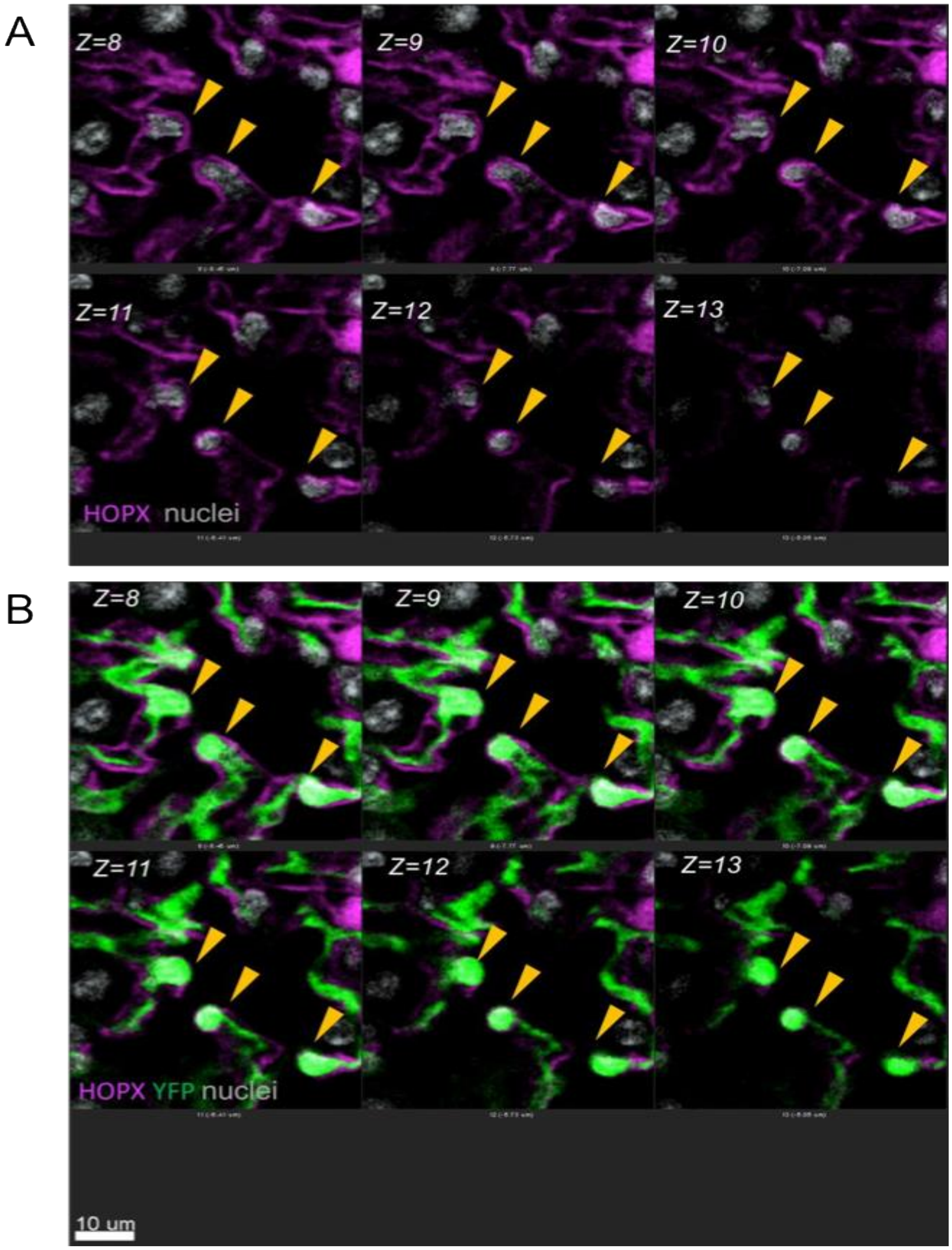

**Supplementary Fig. 7. Tracking of single ATI cells across the z axis within the YFP single cell derived clone.** (A) Chimeric lung sample was stained for expression of HOPX (magenta) across the z-axis of the $YFP^+$ patch and analyzed by confocal microscopy through optical slice z=8 to z=13, clearly detecting single nuclei (gray) in $YFP^+$ cells (B).

A

B

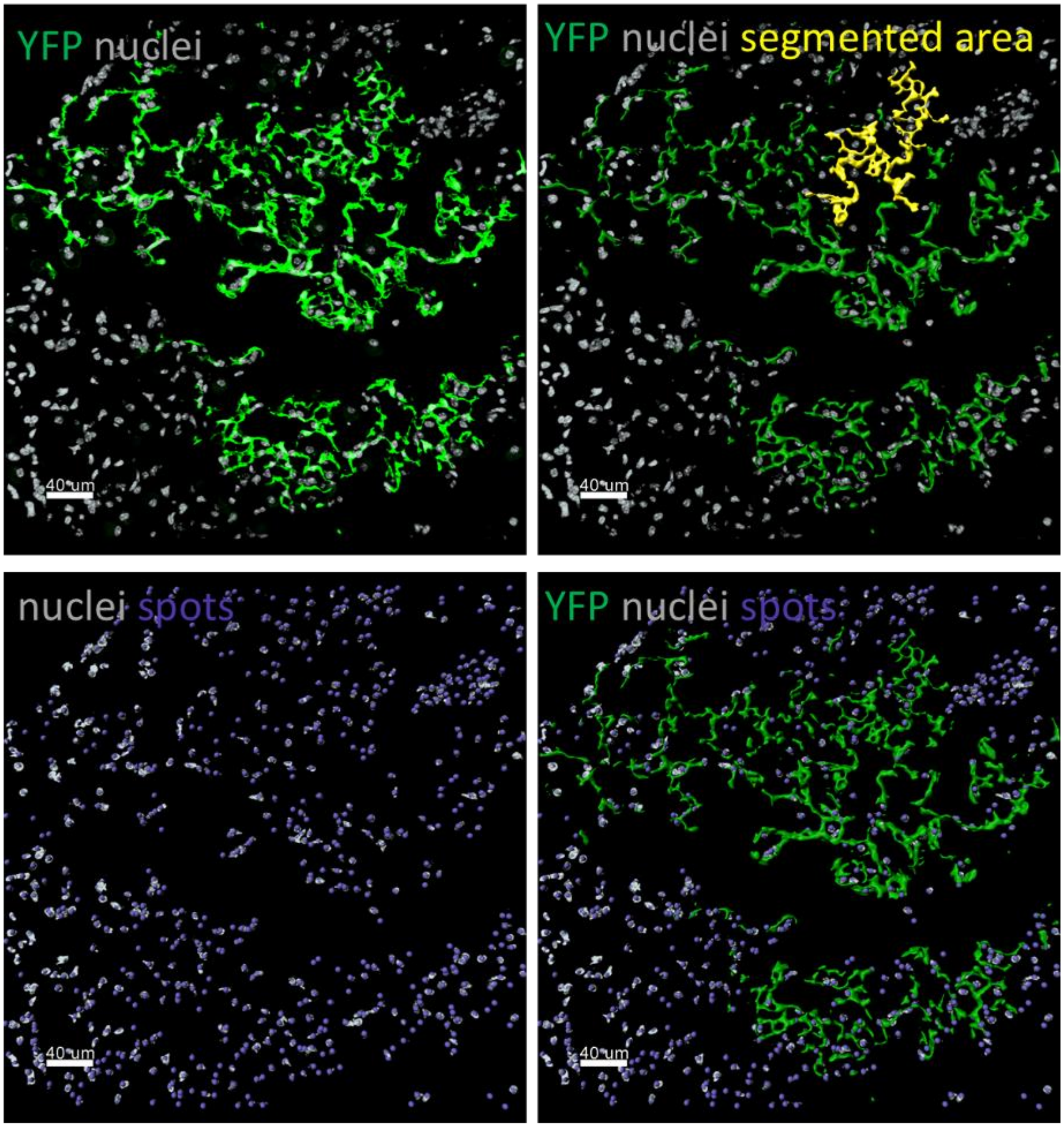

**Supplementary Fig. 8. Calculation of the area and volume of a typical YFP+ patch by segmentation, using the “Surface" Imaris algorithm**. (A) Staining for YFP is shown on the left, and one segment marked in yellow (segment 202, occupying area= 11891.9 $\mu m^2$, and volume is 14286.3 $\mu m^3$) is shown on the right. Details of each segment are shown in Supplementary Table 1. (B) The number of nuclei co-localized in the green channel was determined by using the “spots” function in Imaris. The nuclei in the entire image (left) and in the patch area (right) are shown in grey (counterstaining with Hoechst) and nuclei are marked by blue “spots”. The number of nuclei within the patch was determined manually, to define the average cell area and volume as presented in Table 1 in the main text, and for calculation of the number of nuclei within the donor-derived patch. n=19 patches from 5 mice were evaluated.

| Detailed Area | Unit | Category | Time | ID | Detailed Volume | Unit |
|---|---|---|---|---|---|---|
| 16.3445 | um^2 | Surface | 1 | 140 | 5.32432 | um^3 |
| 40.5082 | um^2 | Surface | 1 | 141 | 22.8096 | um^3 |
| 8.27961 | um^2 | Surface | 1 | 142 | 2.08636 | um^3 |
| 16.3937 | um^2 | Surface | 1 | 143 | 5.53337 | um^3 |
| 17.9543 | um^2 | Surface | 1 | 144 | 6.14308 | um^3 |
| 10.3348 | um^2 | Surface | 1 | 145 | 2.85199 | um^3 |
| 20.5802 | um^2 | Surface | 1 | 146 | 7.20721 | um^3 |
| 32.6675 | um^2 | Surface | 1 | 147 | 12.3155 | um^3 |
| 8.72005 | um^2 | Surface | 1 | 148 | 2.09086 | um^3 |
| 70.8331 | um^2 | Surface | 1 | 149 | 48.1609 | um^3 |
| 77.983 | um^2 | Surface | 1 | 150 | 42.5931 | um^3 |
| 5.96345 | um^2 | Surface | 1 | 151 | 1.09199 | um^3 |
| 7.7703 | um^2 | Surface | 1 | 152 | 1.7203 | um^3 |
| 143.914 | um^2 | Surface | 1 | 153 | 131.665 | um^3 |
| 8.44625 | um^2 | Surface | 1 | 154 | 1.72738 | um^3 |
| 51.7056 | um^2 | Surface | 1 | 155 | 27.2176 | um^3 |
| 3.9719 | um^2 | Surface | 1 | 156 | 0.60369 | um^3 |
| 32.6569 | um^2 | Surface | 1 | 157 | 16.3197 | um^3 |
| 258.714 | um^2 | Surface | 1 | 158 | 250.537 | um^3 |
| 45.7765 | um^2 | Surface | 1 | 159 | 23.2219 | um^3 |
| 5.00764 | um^2 | Surface | 1 | 160 | 0.84239 | um^3 |
| 11.9797 | um^2 | Surface | 1 | 161 | 3.54211 | um^3 |
| 18.2577 | um^2 | Surface | 1 | 162 | 6.17764 | um^3 |
| 14.2455 | um^2 | Surface | 1 | 163 | 4.31616 | um^3 |
| 36.6087 | um^2 | Surface | 1 | 164 | 16.8937 | um^3 |
| 97.1308 | um^2 | Surface | 1 | 165 | 77.4666 | um^3 |
| 18.4612 | um^2 | Surface | 1 | 166 | 6.56054 | um^3 |
| 56.7813 | um^2 | Surface | 1 | 167 | 35.5724 | um^3 |
| 172.51 | um^2 | Surface | 1 | 168 | 136.566 | um^3 |
| 13.8319 | um^2 | Surface | 1 | 169 | 3.25002 | um^3 |
| 30.063 | um^2 | Surface | 1 | 170 | 12.9036 | um^3 |
| 31.679 | um^2 | Surface | 1 | 171 | 13.4568 | um^3 |
| 232.2 | um^2 | Surface | 1 | 172 | 238.402 | um^3 |
| 4.98696 | um^2 | Surface | 1 | 173 | 0.83762 | um^3 |
| 29.0123 | um^2 | Surface | 1 | 174 | 11.6412 | um^3 |
| 36.4056 | um^2 | Surface | 1 | 175 | 18.9612 | um^3 |
| 119.541 | um^2 | Surface | 1 | 176 | 75.9312 | um^3 |
| 254.112 | um^2 | Surface | 1 | 177 | 194.709 | um^3 |
| 19.9033 | um^2 | Surface | 1 | 178 | 7.61773 | um^3 |
| 83.0096 | um^2 | Surface | 1 | 179 | 59.9652 | um^3 |
| 31.2587 | um^2 | Surface | 1 | 180 | 15.1893 | um^3 |
| 338.474 | um^2 | Surface | 1 | 181 | 303.783 | um^3 |
| 122.91 | um^2 | Surface | 1 | 182 | 67.3573 | um^3 |
| 109.444 | um^2 | Surface | 1 | 183 | 99.2379 | um^3 |
| 59.9243 | um^2 | Surface | 1 | 184 | 23.8682 | um^3 |
| 26.1106 | um^2 | Surface | 1 | 185 | 11.5366 | um^3 |
| 759.711 | um^2 | Surface | 1 | 186 | 736.557 | um^3 |
| 252.634 | um^2 | Surface | 1 | 187 | 199.048 | um^3 |
| 688.563 | um^2 | Surface | 1 | 188 | 834.625 | um^3 |
| 31.937 | um^2 | Surface | 1 | 189 | 15.0523 | um^3 |
| 416.495 | um^2 | Surface | 1 | 190 | 364.376 | um^3 |
| 112.43 | um^2 | Surface | 1 | 191 | 91.8863 | um^3 |
| 99.2387 | um^2 | Surface | 1 | 192 | 68.3051 | um^3 |
| 259.456 | um^2 | Surface | 1 | 193 | 219.236 | um^3 |
| 145.738 | um^2 | Surface | 1 | 194 | 114.22 | um^3 |
| 499.129 | um^2 | Surface | 1 | 195 | 440.276 | um^3 |
| 586.565 | um^2 | Surface | 1 | 196 | 655.251 | um^3 |
| 306.581 | um^2 | Surface | 1 | 197 | 287.717 | um^3 |
| 135.404 | um^2 | Surface | 1 | 198 | 99.5614 | um^3 |
| 75.2952 | um^2 | Surface | 1 | 199 | 52.666 | um^3 |
| 406.249 | um^2 | Surface | 1 | 200 | 436.427 | um^3 |
| 38.3077 | um^2 | Surface | 1 | 201 | 20.3177 | um^3 |
| 11891.9 | um^2 | Surface | 1 | 202 | 14286.3 | um^3 |

| Detailed Area | Unit | Category | Time | ID | Detailed Volume | Unit |
|---|---|---|---|---|---|---|
| 11891.9 | um^2 | Surface | 1 | 202 | 14286.3 | um^3 |
| 963.833 | um^2 | Surface | 1 | 203 | 1268.65 | um^3 |
| 379.231 | um^2 | Surface | 1 | 204 | 469.297 | um^3 |
| 68.9838 | um^2 | Surface | 1 | 205 | 48.7618 | um^3 |
| 190.293 | um^2 | Surface | 1 | 206 | 148.752 | um^3 |
| 616.881 | um^2 | Surface | 1 | 207 | 741.694 | um^3 |
| 1263.48 | um^2 | Surface | 1 | 208 | 1450.14 | um^3 |
| 669.835 | um^2 | Surface | 1 | 209 | 883.311 | um^3 |
| 676.85 | um^2 | Surface | 1 | 210 | 869.396 | um^3 |
| 348.75 | um^2 | Surface | 1 | 211 | 304.573 | um^3 |
| 45.9394 | um^2 | Surface | 1 | 212 | 23.2695 | um^3 |
| 476.815 | um^2 | Surface | 1 | 213 | 576.22 | um^3 |
| 704.204 | um^2 | Surface | 1 | 214 | 770.194 | um^3 |
| 644.596 | um^2 | Surface | 1 | 215 | 713.077 | um^3 |
| 129.365 | um^2 | Surface | 1 | 216 | 105.38 | um^3 |
| 125.782 | um^2 | Surface | 1 | 217 | 95.71 | um^3 |
| 952.574 | um^2 | Surface | 1 | 218 | 1064.97 | um^3 |
| 1077.68 | um^2 | Surface | 1 | 219 | 1138.22 | um^3 |
| 105.745 | um^2 | Surface | 1 | 220 | 88.0267 | um^3 |
| 315.668 | um^2 | Surface | 1 | 221 | 316.429 | um^3 |
| 16.7607 | um^2 | Surface | 1 | 222 | 5.71066 | um^3 |
| 22.6827 | um^2 | Surface | 1 | 223 | 9.13971 | um^3 |
| 1488.86 | um^2 | Surface | 1 | 224 | 1597.52 | um^3 |
| 2119.12 | um^2 | Surface | 1 | 225 | 2685.18 | um^3 |
| 12286.6 | um^2 | Surface | 1 | 226 | 16916.6 | um^3 |
| 130.43 | um^2 | Surface | 1 | 227 | 87.5663 | um^3 |
| 203.448 | um^2 | Surface | 1 | 228 | 145.484 | um^3 |
| 433.072 | um^2 | Surface | 1 | 229 | 384.329 | um^3 |
| 799.689 | um^2 | Surface | 1 | 230 | 995.329 | um^3 |
| 7484.37 | um^2 | Surface | 1 | 231 | 10060.8 | um^3 |
| 23.9861 | um^2 | Surface | 1 | 232 | 9.68814 | um^3 |
| 8.17485 | um^2 | Surface | 1 | 233 | 1.7516 | um^3 |
| 1433.78 | um^2 | Surface | 1 | 234 | 1260.7 | um^3 |
| 408.718 | um^2 | Surface | 1 | 235 | 562.232 | um^3 |
| 353.262 | um^2 | Surface | 1 | 236 | 429.672 | um^3 |
| 78.4492 | um^2 | Surface | 1 | 237 | 52.4925 | um^3 |
| 246.498 | um^2 | Surface | 1 | 238 | 248.887 | um^3 |
| 8433.44 | um^2 | Surface | 1 | 239 | 9297.28 | um^3 |
| 1575.21 | um^2 | Surface | 1 | 240 | 1708.32 | um^3 |
| 550.62 | um^2 | Surface | 1 | 241 | 676.285 | um^3 |
| 2266.83 | um^2 | Surface | 1 | 242 | 2463.5 | um^3 |
| 405.486 | um^2 | Surface | 1 | 243 | 421.15 | um^3 |
| 173.503 | um^2 | Surface | 1 | 244 | 116.907 | um^3 |
| 2696.78 | um^2 | Surface | 1 | 245 | 3365.72 | um^3 |
| 135.359 | um^2 | Surface | 1 | 246 | 100.244 | um^3 |
| 183.634 | um^2 | Surface | 1 | 247 | 164.339 | um^3 |
| 299.972 | um^2 | Surface | 1 | 248 | 255.272 | um^3 |
| 373.253 | um^2 | Surface | 1 | 249 | 304.908 | um^3 |
| 594.475 | um^2 | Surface | 1 | 250 | 614.167 | um^3 |
| 182.819 | um^2 | Surface | 1 | 251 | 106.262 | um^3 |
| 299.809 | um^2 | Surface | 1 | 252 | 260.653 | um^3 |
| 36.8396 | um^2 | Surface | 1 | 253 | 18.1531 | um^3 |
| 15.7019 | um^2 | Surface | 1 | 254 | 4.23973 | um^3 |
| 3869.2 | um^2 | Surface | 1 | 255 | 4261.04 | um^3 |
| 175.963 | um^2 | Surface | 1 | 256 | 148.769 | um^3 |
| 123.069 | um^2 | Surface | 1 | 257 | 98.1838 | um^3 |
| 37.8814 | um^2 | Surface | 1 | 258 | 18.1569 | um^3 |
| 28.6989 | um^2 | Surface | 1 | 259 | 10.1116 | um^3 |
| 93.9538 | um^2 | Surface | 1 | 260 | 68.1734 | um^3 |
| 195.275 | um^2 | Surface | 1 | 261 | 166.185 | um^3 |
| 106.259 | um^2 | Surface | 1 | 262 | 61.0401 | um^3 |
| 636.252 | um^2 | Surface | 1 | 263 | 658.788 | um^3 |
| 380.822 | um^2 | Surface | 1 | 264 | 358.194 | um^3 |
| 634.975 | um^2 | Surface | 1 | 265 | 567.781 | um^3 |
| 1294.05 | um^2 | Surface | 1 | 266 | 1300.21 | um^3 |
| **82654** | **um^2** | | | | **95103** | **um^3** |
| total area | | | | | total volume | |

**Supplementary Table 1. Example of monochromatic patch, showed above, area and volume calculation.**

Area (82653.54 μm$^2$) and volume (95102.8 μm$^3$) of a typical YFP+ patch estimated by Imaris software analysis dividing the patch into 266 segments. The patch was found to comprise 73 donor-derived cells, based on determination of nuclei that were colocalized with YFP+ staining (Supplementary Fig. 8). Marking of one typical segment (number 202) is illustrated in yellow in Supplementary Fig. 8.

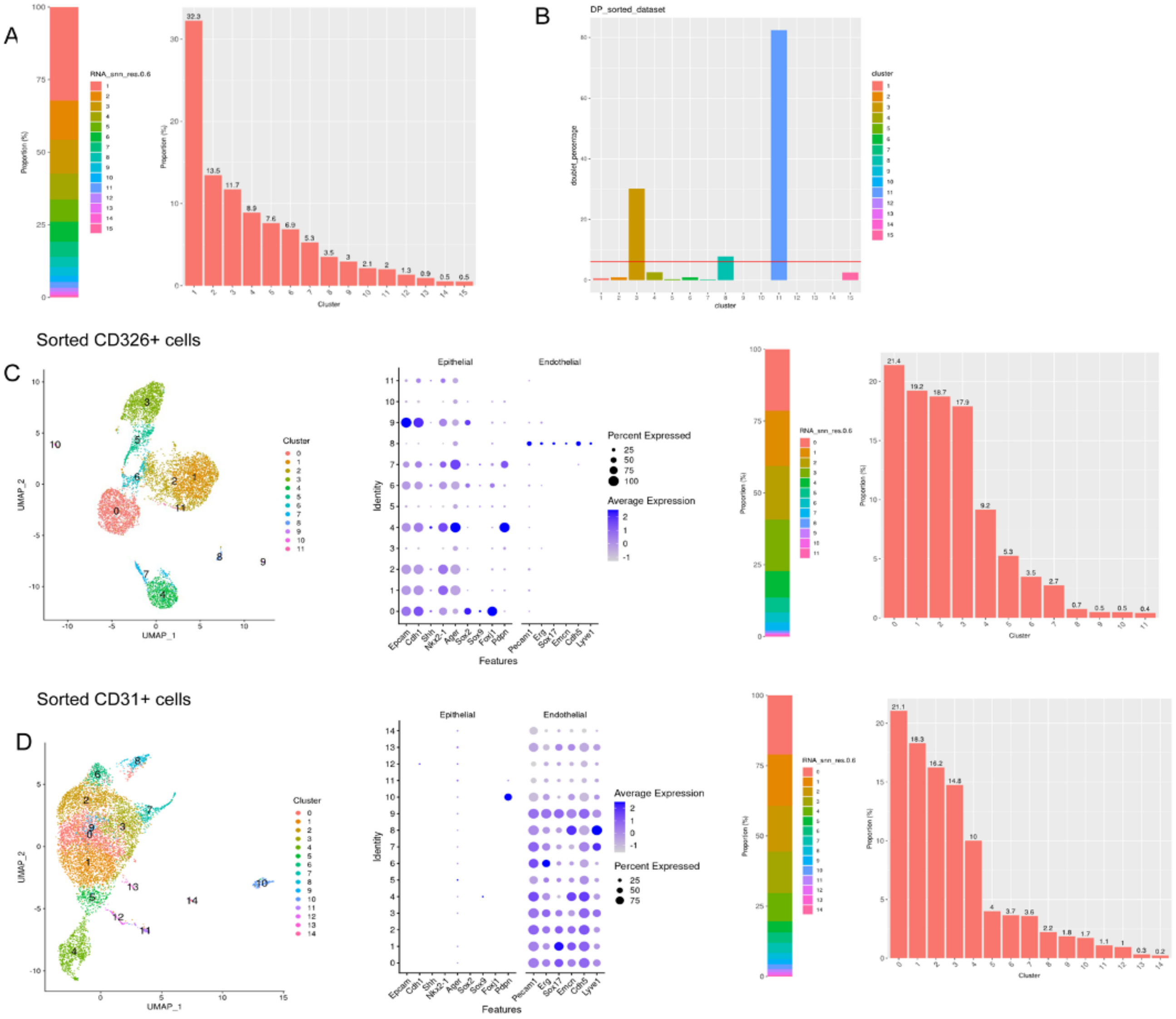

**Supplementary Fig. 9. Transcriptome analysis of FACS purified CD326+CD31+, CD326+ CD31-, CD326- CD31+ cells**

(A) Proportion of the clusters identified at 0.6 resolution within FACS purified double-positive cells (CD326+CD31+), *n*=7689. (B) The expected doublet rate of the double-positive dataset (6.1%) was determined based on the number of recovered cells of this dataset (*n*=8138 before QC steps). Clusters 3, 8, 11 had high doublet percentages, while all other clusters exhibited lower doublet rates, including cluster 5, which maintained a low rate of predicted doublets. (C) UMAP and dot plots of FACS purified epithelial (CD326+ CD31-) cells, (*n*=8151). Expression of epithelial and endothelial genes in different clusters within this population at 0.6 resolution, demonstrating expression of canonical epithelial genes in most of the clusters. The right panel shows the percentage of different clusters within the FACS-purified Epcam+ cells. (D) UMAP and dot plots of FACS purified endothelial (CD326-CD31+) cells, *n*=8703. Expression of epithelial and endothelial genes in different clusters within this population at 0.6 resolution, demonstrating expression of canonical endothelial genes in most of the clusters. The right panel shows the percentage of different clusters within the FACS-purified Pecam1+ cells.

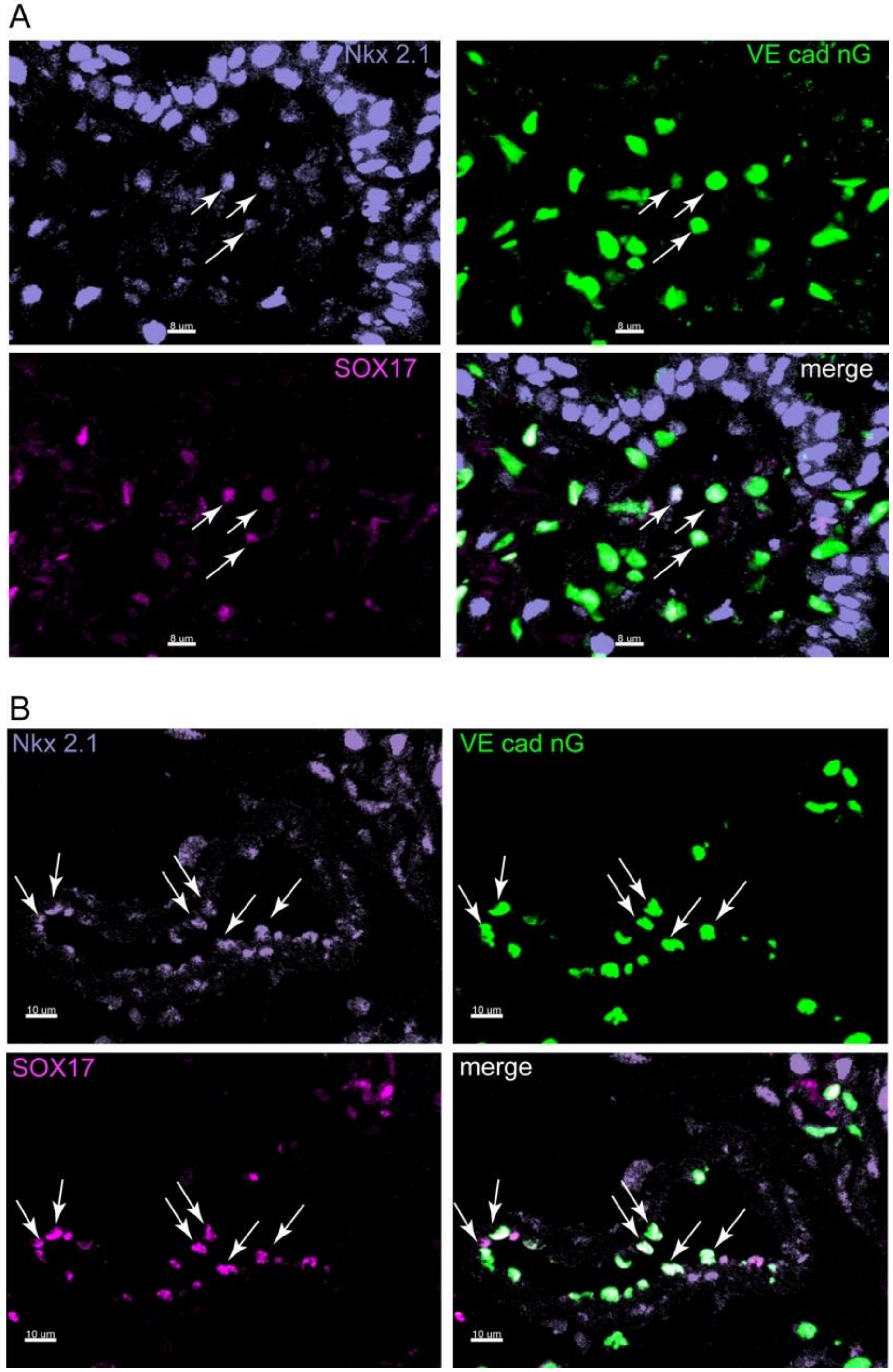

**Supplementary Fig. 10. Detection of lung cells exhibiting markers of both epithelial and endothelial markers.** Immune histological staining of lung tissue from VEcad-Cre-nTnG mouse showing GFP+ cells stained also for NKX 2.1 (violet) and SOX17 (magenta). Triple positive cells are marked by arrows. Scale bar=8um in (A) and 10um (B).

**Supplementary Table 5. Antibodies used in our study.**

| **Primary antibodies** | **Application** | **Catalog No.** | **Dilution** |
|---|---|---|---|
| Rabbit anti- ERG (Abcam) | IHC | AB-92513 | 1:100 |
| Rabbit anti- Nkx2.1 (Abcam) | IHC | AB-76013 | 1:100 |
| Rabbit anti- wide spectrum cytokeratin (Dako) | IHC | Z0622 | 1:100 |
| Rabbit anti- surfactant protein C (Santa-Cruz) | IHC | Sc-13979 | 1:100 |
| Rabbit anti- Aquaporin5 (Millipore) | IHC | CALBIOCHEM 178615 | 1:150 |
| Rat anti-mouse E-cadherin ECCD-2 (Invitrogen) | IHC | 13-1900 | 1:100 |
| Goat anti-mouse CD31 (RD SYSTEMS) | IHC | AF 3628 | 1:100 |
| Rabbit anti SOX17 (Abcam) | IHC | Ab 224637 | 1:100 |
| Guinea pig anti-mouse LAMP3 SYSY (Synaptic Systems) | IHC | 391005 | 1:1000 |
| Rat anti- mouse CD31(Dianova) | IHC | DIA-310-M Clone SZ31 | 1:50-100 |
| Rabbit anti-HOPX | IHC | **HPA030180** | 1:100 |
| Anti- mouse Sca APC-Cy7 (Biolegend) | FACS | 108126 | 1ul/$10^6$ cells |
| Anti- mouse Sca Pacific Blue (Biolegend) | FACS | 108120 | 1ul/$10^6$ cells |

| | | | |
|---|---|---|---|
| Anti- mouse Sca APC (Biolegend) | FACS | 108112 | 1ul/$10^6$ cells |
| Anti- mouse Sca Brilliant Violet 711 (Biolegend) | FACS | 108131 | 1ul/$10^6$ cells |
| Anti- mouse CD45 APC-Cy7 (Biolegend) | FACS | 103116 | 1ul/$10^6$ cells |
| Anti- mouse CD45 PE (Biolegend) | FACS | 103106 | 1ul/$10^6$ cells |
| Anti- mouse CD31 APC (Biolegend) | FACS | 102510 | 1ul/$10^6$ cells |
| Anti- mouse CD31 PE-Cy7 (Biolegend) | FACS | 102418 | 1ul/$10^6$ cells |
| Anti- mouse CD31 PE (Biolegend) | FACS | 102408 | 1ul/$10^6$ cells |
| Anti- mouse CD31 FITC (Biolegend) | FACS | 102506 | 1ul/$10^6$ cells |
| Anti- mouse CD31 PERCP-CY5.5 (Biolegend) | FACS | 102522 | 1ul/$10^6$ cells |
| Anti- mouse CD326 Percp-Cy5.5 (Ep-CAM) (Biolegend) | FACS | 118220 | 1ul/$10^6$ cells |
| Anti- mouse CD326 APC-Cy7 (Ep-CAM) (Biolegend) | FACS | 118218 | 1ul/$10^6$ cells |

| | | | |
|---|---|---|---|
| Anti- mouse CD326 APC (Ep-CAM) (Biolegend) | FACS | 118213 | 1ul/$10^6$ cells |
| Anti- mouse CD326 FITC (Ep-CAM) (Biolegend) | FACS | 118207 | 1ul/$10^6$ cells |
| Anti- mouse CD326 PE (Ep-CAM) (Biolegend) | FACS | 118205 | 1ul/$10^6$ cells |
| Anti-mouse TER-119/Erythroid Cell Pacific Blue (Biolegend) | FACS | 116232 | 1ul/$10^6$ cells |
| Anti-mouse TER-119/Erythroid Cell APC-Cy7 (Biolegend) | FACS | 116223 | 1ul/$10^6$ cells |
| Anti-mouse TER-119/Erythroid Cell APC (Biolegend) | FACS | 116212 | 1ul/$10^6$ cells |
| Anti-mouse CD117 (c-kit) PE (Biolegend) | FACS | 105808 | 1ul/$10^6$ cells |
| SYTOX™ Blue Dead Cell Stain | FACS | S34857 | 1uM |
| 7-AAD (BD Pharmingen) | FACS | 51-68981E | 1ul/$10^6$ cells |
| Anti-mouse CD117 (c-kit) PE (Biolegend) | FACS | 105808 | 1ul/$10^6$ cells |
| ***Secondary antibodies*** | | | |

| | | | |
|---|---|---|---|
| Anti- chicken Alexa Fluor 488 | IHC | 703-545-155 | 1:200 |
| Anti- rabbit Rhodamine Red | IHC | 711-295-152 | 1:200 |
| Anti- rabbit Alexa Fluor 647 | IHC | 711-605-152 | 1:200 |
| Anti- rat Alexa Fluor 594 | IHC | 712-585-150 | 1:200 |
| Anti- goat Alexa Fluor 488 | IHC | 705-545-003 | 1:200 |
| Anti- goat Alexa Fluor 594 | IHC | 705-585-003 | 1:200 |
| Anti- goat AMCA | IHC | 705-155-003 | 1:200 |
| Anti- chicken Alexa Fluor 488 | IHC | 703-545-155 | 1:200 |
| Anti- rabbit Rhodamine Red | IHC | 711-295-152 | 1:200 |
| Anti- rabbit Alexa Fluor 647 | IHC | 711-605-152 | 1:200 |
| Anti- rat Alexa Fluor 594 | IHC | 712-585-150 | 1:200 |
| Anti- goat Alexa Fluor 488 | IHC | 705-545-003 | 1:200 |
| Anti- goat Alexa Fluor 594 | IHC | 705-585-003 | 1:200 |
| Anti- goat AMCA | IHC | 705-155-003 | 1:200 |

| | | | |
|---|---|---|---|
| DyLight™ 405 AffiniPure Donkey Anti-Goat IgG (H+L) | IHC | 705-475-003 | 1:100 |
| Alexa Fluor® 647 AffiniPure Donkey Anti-Goat IgG (H+L) | IHC | 705-605-003 | 1:200 |
| DyLight™ 405 AffiniPure Donkey Anti-Guinea Pig IgG (H+L) | IHC | 706-475-003 | 1:200 |
| DyLight™ 405 AffiniPure Donkey Anti-Rabbit IgG (H+L) | IHC | 711-475-152 | 1:100 |
| DyLight™ 405 AffiniPure Donkey Anti-Rat IgG (H+L) | IHC | 712-475-150 | 1:100 |

*****All the secondary antibodies were produced in donkey and were purchased from Jackson ImmunoResearch or Abcam (unless otherwise indicated).

*Detailed information describing the antibodies, their specificity, cross-reactivity, application and isotype controls is available on the manufacturers' websites.

**Supplementary Movies**

**Supplementary Movie 1**

Two-photon microscopy of whole mount adult ***R26R-Confetti* mouse** lung after administration of two doses of TMX, prior to harvesting cells for transplantation, showing GFP, RFP, YFP and CFP cells randomly distributed throughout the lung tissue. The tissue did not undergo any processing or staining.

**Supplementary Movie 2**

Light sheet microscopy of a cleared chimeric lung, isolated from a mouse treated with Na+6Gy TBI and transplanted with adult ***R26R-Confetti*** mouse lung cells. The endogenous fluorescence was intensified by staining with anti-RFP antibody. The movie shows an example of the RFP patch:  3D rendering across the *xz*-axis is shown on the left side of the screen, the full dimensions of the patch across *xyz* axes are shown on the right.

**Supplementary Movie 3**

Two-photon microscopy of a freshly isolated whole mount chimeric lung 8 weeks post-transplantation, harvested from a mouse treated with Na+6Gy TBI and transplanted with adult ***R26R-Confetti*** mouse lung cells demonstrating discrete monochromatic GFP, RFP, YFP and CFP patches. All monochromatic patches were taken from the same chimeric lung. The tissue did not undergo any processing or staining.
